# Dysgu: efficient structural variant calling using short or long reads

**DOI:** 10.1101/2021.05.28.446147

**Authors:** Kez Cleal, Duncan M. Baird

## Abstract

Structural variation (SV) plays a fundamental role in genome evolution and can underlie inherited or acquired diseases such as cancer. Long-read sequencing technologies have led to improvements in the characterization of structural variants (SVs), although paired-end sequencing offers better scalability. Here, we present dysgu, which calls SVs or indels using paired-end or long reads. Dysgu detects signals from alignment gaps, discordant and supplementary mappings, and generates consensus contigs, before classifying events using machine learning. Additional SVs are identified by remapping of anomalous sequences. Dysgu outperforms existing state-of-the-art tools using paired-end or long-reads, offering high sensitivity and precision whilst being among the fastest tools to run. We find that combining low coverage paired-end and long-reads is competitive in terms of performance with long-reads at higher coverage values.

## Introduction

Analysis of structural variants (SVs) with whole genome or targeted enrichment sequencing is used in the clinic for diagnosing acquired or inherited genetic diseases (1) and for investigating mechanisms of genomic complexity in cancer and other pathologies (2–6). Sequencing using short paired-end reads (PE) is well established for genomic analysis due to mature workflows and low sequencing costs, although increasingly, long-read (LR) sequencing technologies are being utilized for these purposes. These LR sequencing platforms permit much longer read-lengths which can potentially lead to improvements in mapping to repetitive or complex regions of the reference genome, and advantages for detecting SVs. However, the better scalability of paired-end technologies, with further improvements in development (7), means that SV calling with shorter reads is likely to remain an area of interest.

SVs are usually defined as genomic rearrangement events over an arbitrary size of 50 bp, falling into categories such as deletions (DEL), insertions (INS), duplications (DUP), inversions (INV) or translocations (TRA) (1). SVs below this threshold are often termed indels, although these can sometimes result from more complex events such as duplication, inversion or translocation. These labels are useful in conceptualizing simple genome rearrangements in terms of the reference genome structure, although complex SVs occurring in the germline or during cancer progression, can complicate interpretation.

SVs can be detected in sequencing data using a variety of methods. For PE data, single alignments only span relatively small within-read SVs (indels) due to limited read-length, so information of SVs must be gleaned from assessing discordant mappings, changes in read-depth and the occurrence of split-reads which straddle breaksites (8). Recent methods also employ de novo assembly of SV-derived reads and further rounds of SV discovery through re-mapping of derived contigs to the reference genome (9, 10)28/05/2021 07:05:00. Alignment free methods are also possible, by analysing differences in k-mer content between a sample and reference (11). For LR sequences, SVs up to several kb can be detected within alignments due to the long read-lengths involved, and split-reads, changes in read depth and assembly of SV-reads can be utilized (8).

A large number of bioinformatics tools have been developed for detecting SVs using PE or LR data, although recent benchmarking studies highlight that existing algorithms are often limited in their ability to detect all classes and sizes of SVs, and there is still considerable room for improvement (12–14). The approach of quality filtering of putative SVs also differs widely between tools. In the simplest case variants are filtered based on the weight of evidence or number of supporting reads, although choosing suitable thresholds can be difficult and higher read-depths have also been associated with false positives (13). Statistical methods for quality scoring have been employed, for example the PE caller Manta employs Bayesian inference using read fragments supporting an allele to estimate a likelihood, followed by manual filtering (9). The LR caller nanovar utilizes a neural network classifier trained on simulated datasets, where 14 input features of each putative SV are used to classify events (15). To build on these advances, we considered that performance may be enhanced from training using non-simulated datasets. Additionally, we identified that there is an unmet need for an SV caller capable of analysing both PE and LR datasets.

Here, we present our SV calling software dysgu, which can rapidly call SVs from PE or LR data, across all size categories. Conceptually, dysgu identifies SVs from alignment cigar information as well as discordant and split-read mappings. Dysgu employs a fast consensus sequence algorithm, inspired by the positional de Brujin graph, followed by remapping of anomalous sequences to discover additional small SVs. A machine learning classifier is then employed to generate a useful quality score which can be used to prioritize variants.

## Results

Dysgu is a general purpose *de novo* SV and indel caller that can analyse PE or LR sequencing datasets. SV-associated reads are first identified by assessing alignment gaps, split-read and discordant mappings, soft-clipped reads and read-depth changes. SV signals are clustered on a graph and contigs are generated for putative breakpoints. One-end anchored SVs - events with a single soft-clipped sequence without a corresponding mapping, are re-aligned to the reference genome to identify additional small SVs. Putative SV events are labelled with a rich set of features describing sequencing or mapping error metrics and supporting evidence. Events are further classified using a machine learning model to prioritise variants with higher probability.

### Testing datasets

To assess precision and recall statistics we utilized benchmark datasets provided by the Genome in a Bottle (GIAB) consortium. Primarily, we assesses a germline call set derived from the Ashkenazi son sample (HG002) that combines five sequencing technologies and 68 call sets plus manual curation into a high quality and comprehensive benchmark (16). The HG002 benchmark is stratified into high confidence regions (Tier 1), where precision and recall can be confidently determined, as well as less confident regions (Tier 2, followed by ‘all’ regions) which potentially involve more complex genomic regions, or the completeness of the benchmark is uncertain. However, as only SVs ≥ 50 bp appear in Tier 1 regions, we also analysed all unfiltered SVs in the GIAB dataset which has a minimum SV size threshold ≥ 20 bp, appreciating that the ‘All-regions’ benchmark shows lower completeness compared to Tier 1 regions. In addition, we assessed recall on the HG001 cell line that has corresponding deletion calls (≥ 50 bp) provided by GIAB (17). As the machine-learning classifier that dysgu employs was trained using calls derived from HG001 (see Methods), we did not assess precision using this dataset.

### Performance using paired-end short reads

Dysgu was tested on HG002 at coverages of 20× (Figure 1, Table 1, 2, Supplemental_Table_S1.pdf) and 40× (Supplemental_Fig_S1.pdf, Supplemental_Table_S2.pdf - Supplemental_Table_S4.pdf) using Illumina 148 bp paired-end reads. Performance was compared to the popular SV callers manta (9), delly (18), and lumpy (19). We also compared indel calling performance with strelka (20) and gatk down to a size of 30 bp. Strelka calls indels up to 50 bp whilst gatk calls deletions and insertions to around the insert size.

**Figure 1.**
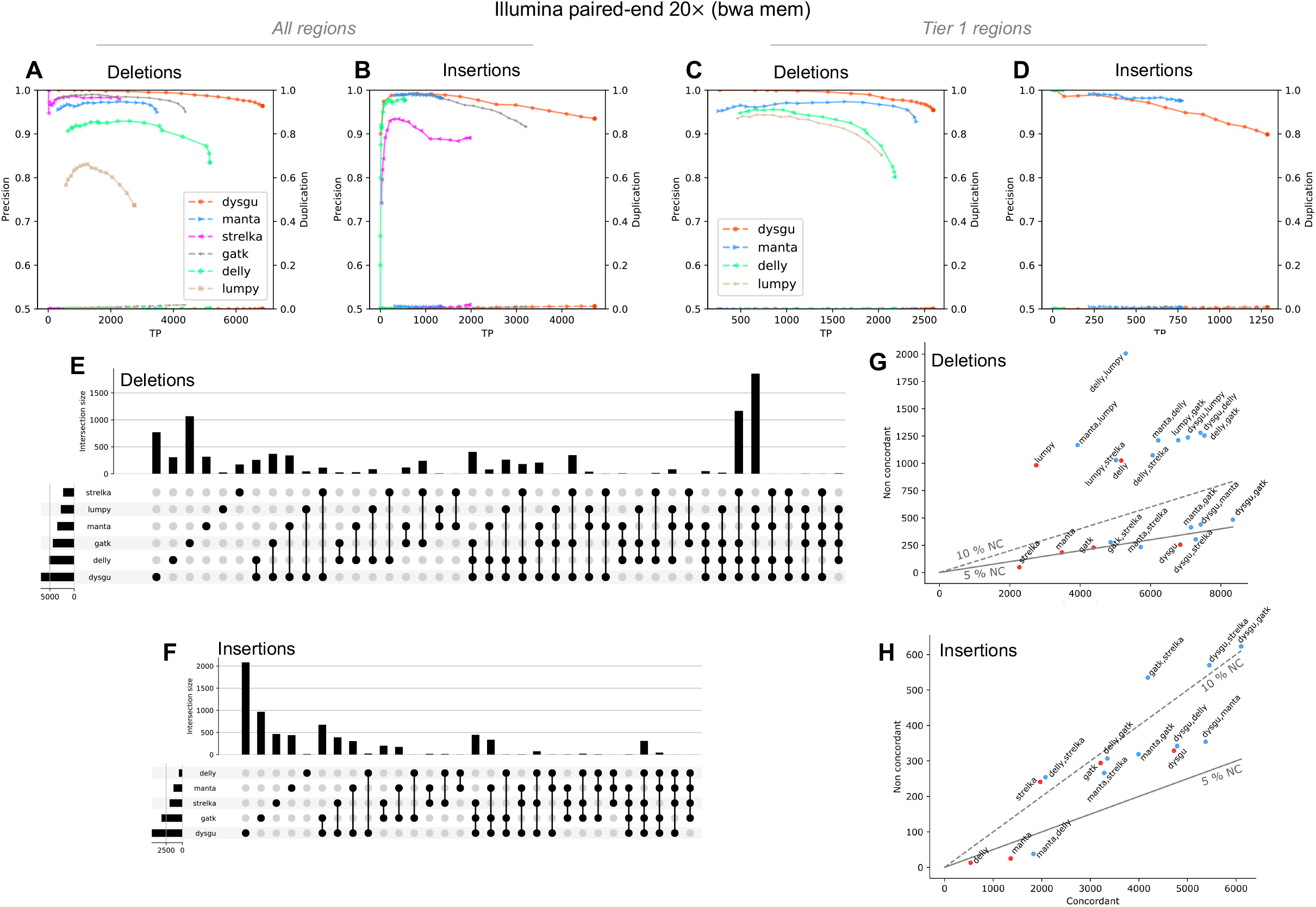
Performance of dysgu using 20× PE reads. Dysgu was compared to SV callers manta, delly and lumpy, and indel callers strelka and gatk, using the HG002 benchmark. Precision-recall curves are shown for all genomic regions (A, B), as well as high-confidence Tier 1 regions (C, D). The secondary y-axis indicates duplicate true-positives (TP) as a fraction of true-positive calls. Intersections and aggregates of intersections of SV calls for the all-regions benchmark are displayed using an upset plot (E, F). To investigate combinations of SV callers, the union of true-positives between callers (labelled concordant), was plotted against the sum of false-positives (labelled non concordant) (G, H). The 5 and 10 % non-concordance (NC) is also illustrated as a solid or dashed line, respectively.

**Table 1.** Performance using PE 20× data on the HG002 ‘Tier 1 regions’ benchmark. The numbers of deletion (DEL) and insertion (INS) variants are quantified. Duplication is defined as the ratio of duplicate true-positive calls to the number of true-positive calls. TP – true-positive, FP – false-positive. Best scores are shaded blue.

|  | TP |  | FP |  | Precision |  | Recall |  | Duplication |  | F1 |  |
| --- | --- | --- | --- | --- | --- | --- | --- | --- | --- | --- | --- | --- |
|  | DEL | INS | DEL | INS | DEL | INS | DEL | INS | DEL | INS | DEL | INS |
| dysgu | 2601 | 1293 | 119 | 134 | 0.956 | 0.906 | 0.617 | 0.238 | 0.001 | 0.012 | 0.750 | 0.377 |
| manta | 2411 | 775 | 187 | 19 | 0.928 | 0.976 | 0.572 | 0.142 | 0.000 | 0.008 | 0.708 | 0.249 |
| delly | 2178 | 58 | 536 | 0 | 0.803 | 1.000 | 0.517 | 0.011 | 0.001 | 0.000 | 0.629 | 0.021 |
| lumpy | 2037 |  | 350 |  | 0.853 |  | 0.483 |  | 0.001 |  | 0.617 |  |

**Table 2.**
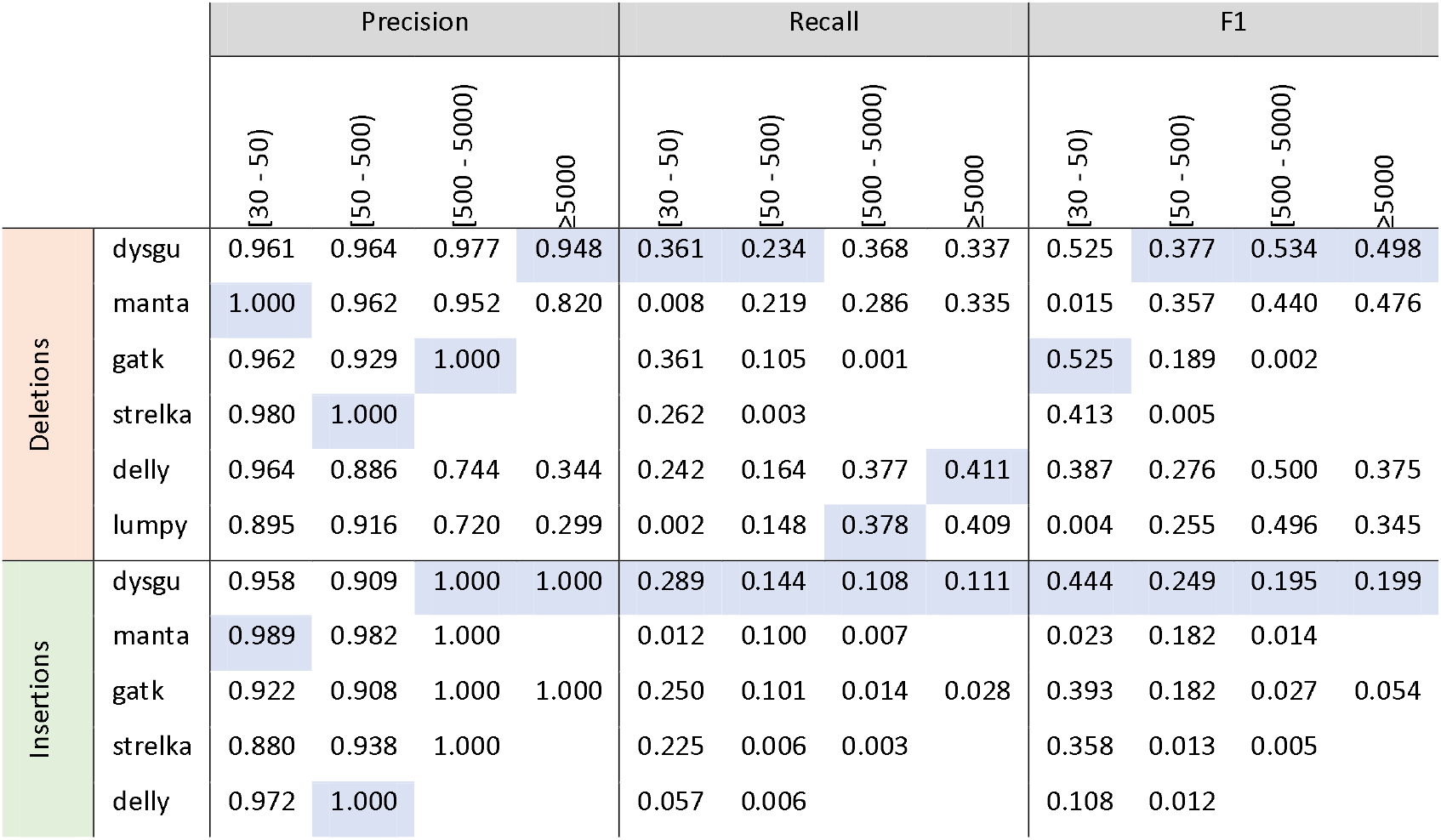
SV calling stratified by size using PE 20× data on the HG002 the ‘All-regions’ benchmark. Best scores are shaded blue.

For Tier 1 SVs at 20× coverage, dysgu called the largest number of true deletions and insertions (n = 3894), with 708 more variants called than the next best caller manta (n = 3186) (Table 1). Precision-recall curves indicated that probability values estimated by dysgu using machine learning were useful for stratifying variants by quality, with higher probability values correlating with precision (Table 1A-D). Dysgu had the highest precision for deletion calls (95.6 %), as well as the highest recall for deletions (61.7 %) and insertions (23.8 %). Manta showed the highest precision for insertion variants (97.6 % vs dysgu 90.6 %) but had a lower recall (14.2 %) than dysgu. As a percentage value, dysgu called 7.9 % more deletions and 67 % more insertions than manta. Overall, dysgu showed higher F1 scores than the next best caller, manta, with an F1 score 4.2 % higher for deletions and 12.8 % higher for insertions. We also assessed the level of duplication, defined as the ratio of duplicated true-positive calls relative to unique true-positive calls. The problem of duplication arises when a single SV event leads to multiple calls in the output file. Generally, all PE callers displayed a low level of duplication below < 1.5 % (Table 1).

We also stratified variants by size using the All-regions benchmark to investigate size constraints of SV calling (Table 2, Supplemental_Table_S4.pdf). For deletions in the 30 – 50 bp range, dysgu showed similar performance to gatk with similar precision, recall and F1 scores. For insertions in the 30 - 50 bp range, dysgu showed higher precision (95.8 %) and recall (28.9 %) than strelka and gatk.

For SVs ≥ 50 bp, dysgu showed a good balance of precision and recall across all size ranges with the highest F1 scores among callers (Table 2). For deletion SVs dysgu generally displayed the highest precision but showed a lower recall for large SVs. For example, delly showed a higher recall than dysgu for deletions ≥ 5000 bp (41.1 % vs 33.7 %), but only had a precision of 34.4 % vs dysgu 94.8 %.

For insertion SVs, dysgu showed the highest recall, but manta displayed the best precision of 98.2 %. Dysgu was the best caller for identifying loci with large insertions (≥ 500 bp) finding n=386, vs manta n=23 and gatk n=49. However, as dysgu utilizes insert size statistics to estimate large insertions length, calculated insertion sizes are expected to be less accurate compared to *de novo* assembly-based callers such as manta and gatk (data not shown).

At 40× coverage, all callers displayed improved recall and F1 scores although at the expense of lower precision (Supplemental_Fig_S1.pdf, Supplemental_Table_S2.pdf - Supplemental_Table_S4.pdf). Interestingly, this phenomenon was also reported in a recent benchmarking study suggesting that at higher coverage values, absolute numbers of sequencing and mapping artifacts are more likely to be mistaken for SV events with low allelic fraction (12). Overall, at 40× coverage dysgu maintained a good balance of precision and recall compared to other callers, in line with 20× coverage, showing the highest F1 score for deletions and insertion calls.

We next investigated the intersection of variant calls between tools, or the set of SVs shared between tools, and displayed results using an upset plot (Figure 1E, F), which quantifies the sizes of SV call sets, their intersections, and aggregates of intersections (21). Assessing Tier 1 SVs in the HG002 benchmark, dysgu showed the largest number of unique calls (both deletions n=154, and insertions n=815) followed by manta (n=135 deletions, n=295 insertions). Including indel callers and analysing all SVs changed the conclusion slightly. In this case, gatk found the most unique deletions events (n=1928, vs dysgu n=622) and the second highest number of unique insertion events (n=1610 after dysgu n=1800).

Recent studies have investigated combining the output of different SV callers to boost performance (22–24). To gauge the performance of different combinations of callers we assessed the union of true positive calls (labelled as concordant) and compare with the sum of false positives (labelled non-concordant) as a proxy for the false positive rate (Figure 1G, H). The best combination of callers using the All-regions benchmark appeared to be dysgu and gatk which together found 3069 deletions and 4368 insertions absent from other callers.

We additionally tested the recall of tools against the HG001 deletion call set, comparing unfiltered variants for all callers. Dysgu demonstrated the highest recall (93.61%), followed by manta (89.84 %), delly (84.38 %) and lumpy (81.61 %).

To summarise, using PE data, dysgu was generally the most performant tool showing a good balance of precision and recall across SV types and size ranges.

### Performance using long reads

We tested dysgu against the HG002 benchmark using PacBio HiFi reads at approximately 8× (Figure 2, Tables 3-4, Supplemental_Fig_S2.pdf, Supplemental_Table_S5.pdf - Supplemental_Table_S8.pdf) and 15× coverage (Supplemental_Fig_S3.pdf, Supplemental_Table_S9.pdf - Supplemental_Table_14.pdf), and using Oxford nanopore reads at 13× coverage (Supplemental_Fig_S4.pdf, Supplemental_Table_S15.pdf - Supplemental_Table_S20.pdf). Performance was compared against recently published LR callers nanovar (15), sniffles (25) and svim (26), using reads aligned by minimap2 (27) (Figures 2, Table 3 - 4), or ngmlr (25) (Supplemental_Fig_S2.pdf, Supplemental_Table_S5.pdf). Aligning reads using ngmlr tended to give slightly higher precision among all SV callers although F1 scores were also slightly reduced, particularly for insertion variants (Supplemental_Table_S5.pdf - Supplemental_Table_S7.pdf).

**Figure 2.**
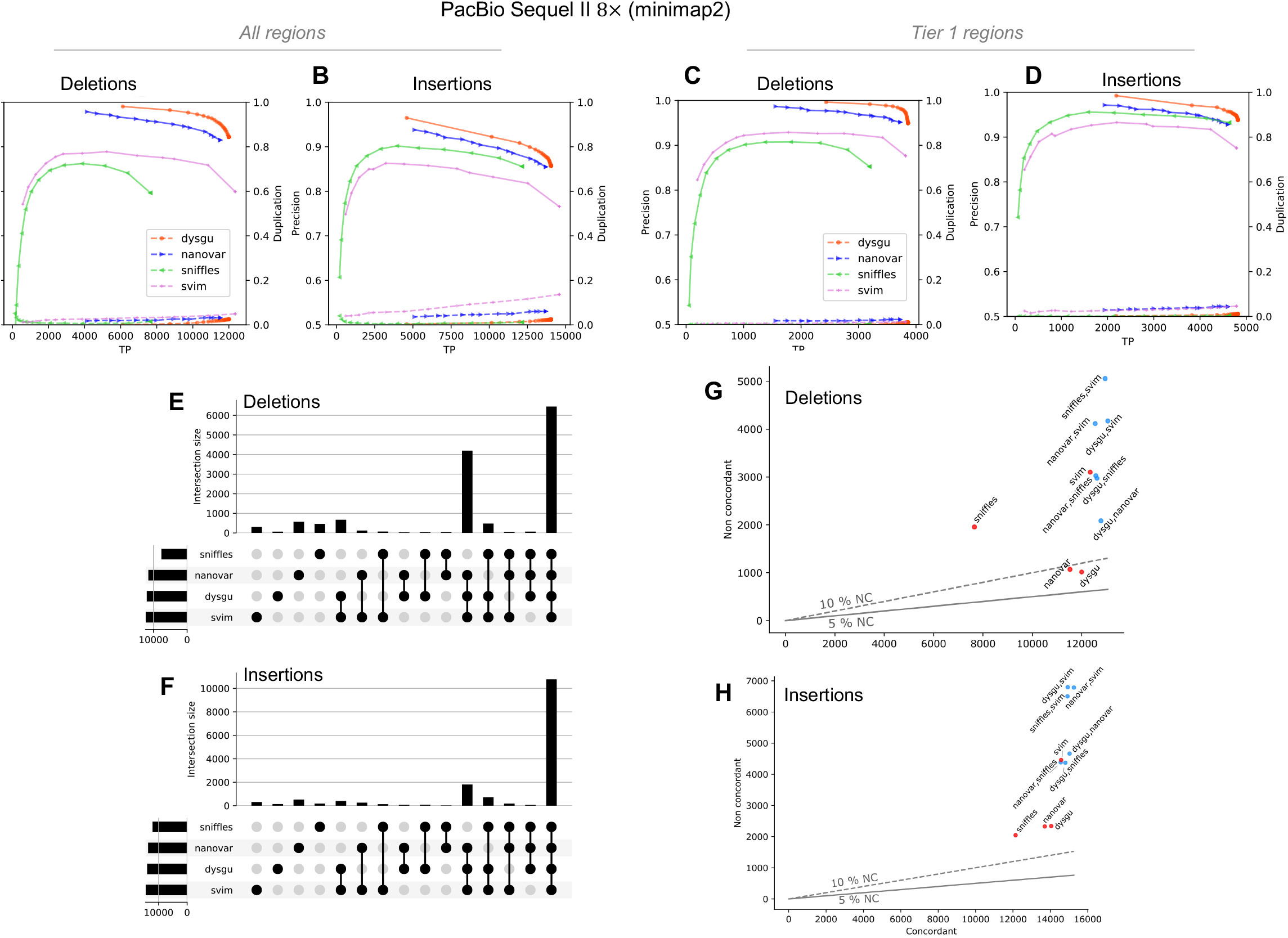
Performance of dysgu using PacBio reads. Precision-recall curves are shown for all genomic regions (A, B), as well as high-confidence Tier 1 regions (C, D). Analysis of SV intersections and aggregates of intersections for the all-regions benchmark are displayed using an upset plot (E, F). The combinations of SV callers was assessed by plotting the union of true-positives (labelled concordant), against the sum of false-positives (labelled non concordant) (G, H). The 5 and 10 % non-concordance (NC) are shown as a solid or dashed line, respectively.

**Table 3.**
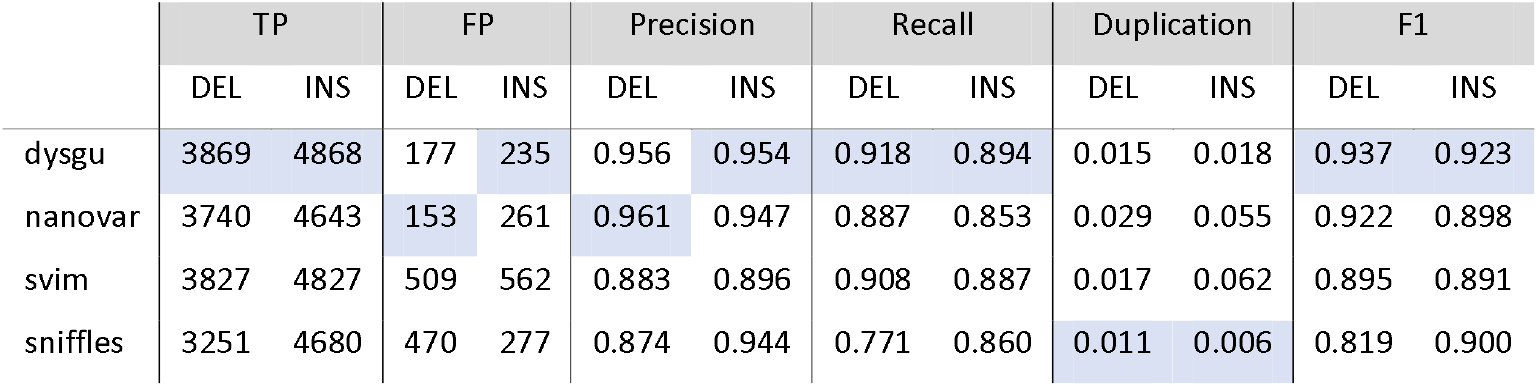
Performance using PacBio Sequel II reads at 8× coverage on HG002 Tier 1 regions. Duplication is defined as the ratio of duplicate true-positive calls to the number of true-positive calls. TP – true-positive, FP – false-positive. Best scores are shaded blue.

|  | TP |  | FP |  | Precision |  | Recall |  | Duplication |  | F1 |  |
| --- | --- | --- | --- | --- | --- | --- | --- | --- | --- | --- | --- | --- |
|  | DEL | INS | DEL | INS | DEL | INS | DEL | INS | DEL | INS | DEL | INS |
| dysgu | 3869 | 4868 | 177 | 235 | 0.956 | 0.954 | 0.918 | 0.894 | 0.015 | 0.018 | 0.937 | 0.923 |
| nanovar | 3740 | 4643 | 153 | 261 | 0.961 | 0.947 | 0.887 | 0.853 | 0.029 | 0.055 | 0.922 | 0.898 |
| svim | 3827 | 4827 | 509 | 562 | 0.883 | 0.896 | 0.908 | 0.887 | 0.017 | 0.062 | 0.895 | 0.891 |
| sniffles | 3251 | 4680 | 470 | 277 | 0.874 | 0.944 | 0.771 | 0.860 | 0.011 | 0.006 | 0.819 | 0.900 |

**Table 4.** Long-read performance as a function of SV size. PacBio Sequel II reads at 8× coverage were assessed using the HG002 ‘all-regions’ benchmark. Best scores are shaded blue.

|  |  | Precision |  |  |  | Recall |  |  |  | F1 |  |  |  |
| --- | --- | --- | --- | --- | --- | --- | --- | --- | --- | --- | --- | --- | --- |
|  |  | [30, 50) | [50, 500) | [500, 5000) | ≥5000 | [30, 50) | [50, 500) | [500, 5000) | ≥5000 | [30, 50) | [50, 500) | [500, 5000) | ≥5000 |
| Deletions | dysgu | 0.932 | 0.930 | 0.939 | 0.929 | 0.551 | 0.505 | 0.438 | 0.321 | 0.693 | 0.654 | 0.597 | 0.477 |
|  | nanovar | 0.939 | 0.933 | 0.882 | 0.730 | 0.504 | 0.475 | 0.443 | 0.354 | 0.656 | 0.629 | 0.590 | 0.477 |
|  | svim | 0.850 | 0.786 | 0.835 | 0.925 | 0.565 | 0.519 | 0.434 | 0.276 | 0.679 | 0.625 | 0.571 | 0.425 |
|  | sniffles | 0.920 | 0.875 | 0.636 | 0.411 | 0.261 | 0.362 | 0.440 | 0.354 | 0.407 | 0.512 | 0.520 | 0.380 |
| Insertions | dysgu | 0.829 | 0.861 | 0.946 | 0.887 | 0.566 | 0.589 | 0.509 | 0.249 | 0.672 | 0.699 | 0.662 | 0.389 |
|  | nanovar | 0.843 | 0.874 | 0.887 | 0.364 | 0.531 | 0.569 | 0.506 | 0.126 | 0.651 | 0.690 | 0.644 | 0.188 |
|  | svim | 0.765 | 0.759 | 0.904 | 0.961 | 0.585 | 0.609 | 0.524 | 0.194 | 0.663 | 0.676 | 0.664 | 0.322 |
|  | sniffles | 0.854 | 0.864 | 0.877 | 0.852 | 0.452 | 0.541 | 0.469 | 0.182 | 0.591 | 0.665 | 0.611 | 0.300 |

Assessing Tier 1 SVs from the HG002 benchmark, dysgu had the highest recall for deletions (91.8 %) and insertions (89.4 %) and the highest precision for insertion calls (95.4 %). Dysgu also had the highest F1 score for deletions (0.937) and insertions (0.923) but was closely followed by nanovar with F1 scores of 0.922 and 0.898 for deletions and insertions, respectively (Figure 2 and Table 3).

Expanding the testing set to all regions and a minimum size of 30 bp, svim showed the highest recall (0.334 for deletions and 0.403 for insertions) (Supplemental_Table_S6.pdf - Supplemental_Table_S7.pdf). Dysgu and nanovar displayed similar precision scores, but overall dysgu displayed the highest F1 scores (0.482 for deletions and 0.537 for insertions) (Supplemental_Table_S6.pdf). Svim showed marginally lower F1 scores (0.475 for deletions and 0.534 for insertions), although we noticed that svim showed a higher level of duplication. Additionally, for some callers this problem was more acute when analysing Oxford nanopore reads, with for example, svim showing a duplication ratio of 0.58 for insertion calls in Tier 1 regions (Supplemental_Figure_S4.pdf, Supplemental_Table_S15.pdf). Among callers, sniffles and dysgu generally showed the lowest duplication rates, although dysgu had a consistently higher recall.

Analysing the intersection of SVs, we found that most callers seemed to identify similar sets of SVs indicating that combining SV callers might only lead to small gains in sensitivity (Figure 2E-H).

Similar to Illumina data, increasing the coverage of PacBio HiFi data increased the recall of SV callers and F1 scores, but at the expense of reduced precision. At 15× coverage, dysgu had the highest F1 scores for deletions and insertions for Tier 1, whilst showing a low level of duplication (Supplemental_Table_S9.pdf). Sensitivity of SV detection was also assed using the HG001 deletion benchmark (≥ 50 bp in size). Using PacBio reads at 5× coverage dysgu showed the highest recall (77.35 %) compared to other callers (nanovar 75.97, sniffles 70.52, svim 73.73 %). Likewise, dysgu showed the highest recall using 13× ONT reads (96.41 %) compared to other callers (nanovar 91.67, sniffles 95.89, svim 95.25 %).

In summary, dysgu demonstrated a high level of performance of LR datasets, with generally the best balance of precision and recall across SV sizes and categories.

### Combining short and long reads for improved performance

Dysgu supports merging of SVs from different runs using a ‘merge’ command making it trivial to integrate calls from different sequencing technologies. After merging, additional tags are added to the output file corresponding to the maximum and mean probability across samples, with the probability determined by the machine learning classifier.

We used dysgu to assess different combinations of sequencing technology including PacBio (8× and 15×), ONT (13×) and Illumina paired-end reads (20× and 40×), by filtering calls with a maximum model probability ≥ 0.5 for PacBio, or ≥ 0.35 for ONT combinations (Table 5). Testing against the All-regions benchmark, the addition of Illumina reads consistently led to performance improvements when combined with PacBio or ONT, especially for deletion calls (Table 5). The largest increases in recall were seen from adding 40× Illumina calls, although 20× Illumina calls also led to noticeable increases. For example, adding 40× Illumina calls to 8× PacBio calls identified an additional 1010 deletions and 1103 insertions for the All-regions benchmark, or 141 deletions and 85 insertions for Tier 1 regions. F1 scores improved for the All-regions benchmark, increasing by 2.77 % for deletions and 2.57 % for insertions. Surprisingly, combining Illumina calls with PacBio 8×, appeared to be similar in performance to PacBio calls at a higher coverage value 15×.

**Table 5.**
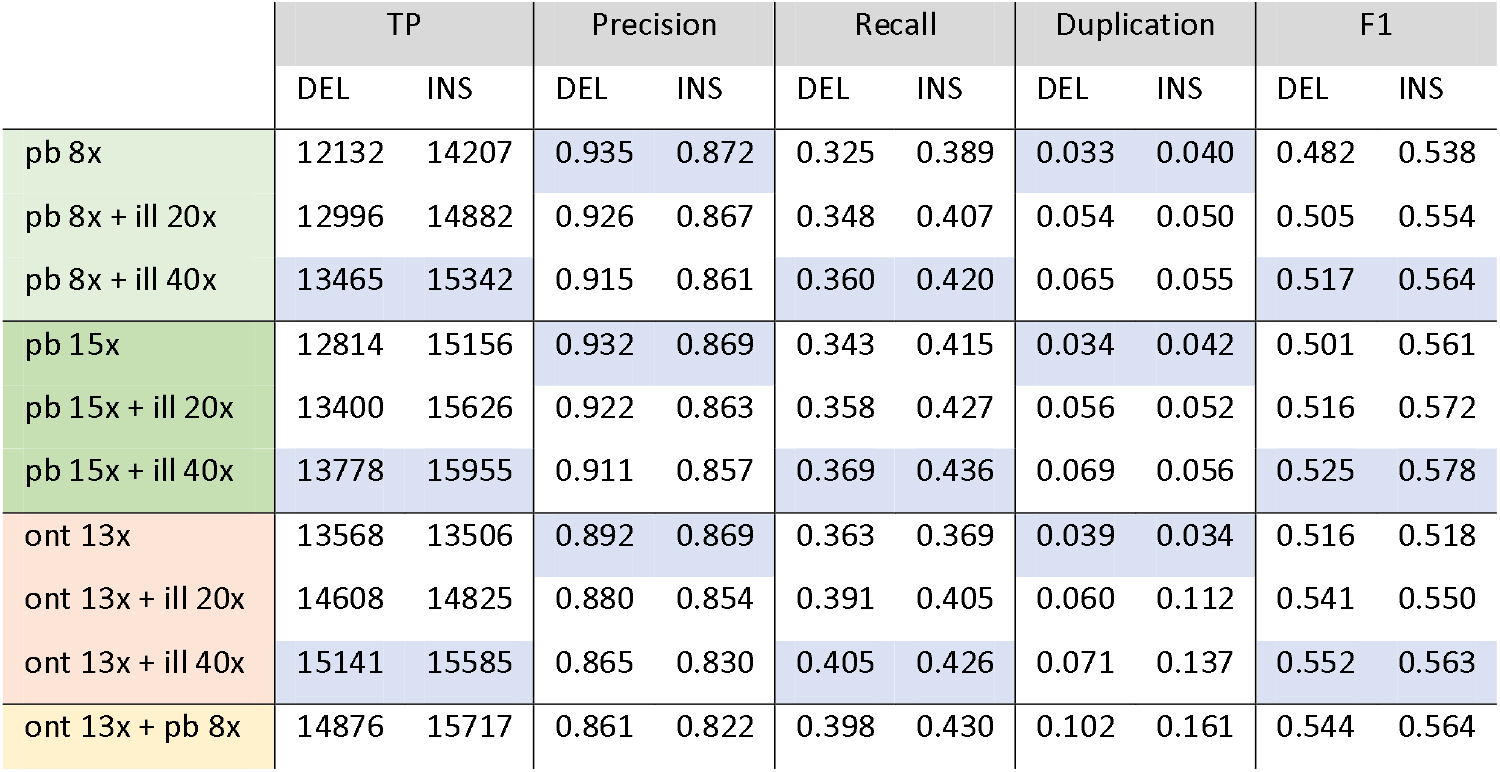
Performance of combinations of sequencing platforms using the HG002 ‘all-regions’ benchmark. pb – PacBio, ill – Illumina, ont – Oxford Nanopore Technologies. Best scores are shaded blue.

|  | TP |  | Precision |  | Recall |  | Duplication |  | F1 |  |
| --- | --- | --- | --- | --- | --- | --- | --- | --- | --- | --- |
|  | DEL | INS | DEL | INS | DEL | INS | DEL | INS | DEL | INS |
| pb 8x | 12132 | 14207 | 0.935 | 0.872 | 0.325 | 0.389 | 0.033 | 0.040 | 0.482 | 0.538 |
| pb 8x + ill 20x | 12996 | 14882 | 0.926 | 0.867 | 0.348 | 0.407 | 0.054 | 0.050 | 0.505 | 0.554 |
| pb 8x + ill 40x | 13465 | 15342 | 0.915 | 0.861 | 0.360 | 0.420 | 0.065 | 0.055 | 0.517 | 0.564 |
| pb 15x | 12814 | 15156 | 0.932 | 0.869 | 0.343 | 0.415 | 0.034 | 0.042 | 0.501 | 0.561 |
| pb 15x + ill 20x | 13400 | 15626 | 0.922 | 0.863 | 0.358 | 0.427 | 0.056 | 0.052 | 0.516 | 0.572 |
| pb 15x + ill 40x | 13778 | 15955 | 0.911 | 0.857 | 0.369 | 0.436 | 0.069 | 0.056 | 0.525 | 0.578 |
| ont 13x | 13568 | 13506 | 0.892 | 0.869 | 0.363 | 0.369 | 0.039 | 0.034 | 0.516 | 0.518 |
| ont 13x + ill 20x | 14608 | 14825 | 0.880 | 0.854 | 0.391 | 0.405 | 0.060 | 0.112 | 0.541 | 0.550 |
| ont 13x + ill 40x | 15141 | 15585 | 0.865 | 0.830 | 0.405 | 0.426 | 0.071 | 0.137 | 0.552 | 0.563 |
| ont 13x + pb 8x | 14876 | 15717 | 0.861 | 0.822 | 0.398 | 0.430 | 0.102 | 0.161 | 0.544 | 0.564 |

However, Tier 1 regions generally did not show increased F1 scores despite increased recall, which was caused by an inflation of the false-positives rate (Supplemental_Table_S21.pdf). Additionally, we assessed Tier 1+2 regions which include more complicated genomic loci than Tier 1. Tier 1+2 regions also showed improved F1 scores, with 8× PacBio + 40× Illumina F1 scores increasing by 3.0 points for deletions and 1.9 for insertions (Supplemental_Table_S22.pdf). We speculate that Illumina data may enhance SV calling at complicated genomic regions that are not trivial to map for LR mappers. Additionally, PE data may help fill-in the gaps for LR datasets in regions of low or zero coverage.

Combining sequencing technologies for improved SV discovery has not received much attention, although with the increasing prevalence of LR sequencing, and other non-standard techniques such as linked-read or HiC, we suggest that this would be an interesting avenue for future research.

### Runtime

We tested runtime using an Intel(R) Xeon(R) CPU E5-2680 v3 @ 2.50GHz Linux machine with 256 GB of system memory. For Ilumina data, dysgu was the fastest tool using a single-core, analysing 40× coverage data in 75 mins and using 5.6 GB memory (Table 6), which was almost twice as quick as the next fastest tool, delly. Manta was 4.85 times slower than dysgu to run on a single core, but used the least memory (0.244), and can also be run in parallel efficiently (data not shown). For PacBio HiFi reads analysed on a single core, dysgu was the second fastest tool after svim, analysing 8× coverage sample in 8 mins and using 0.35 GB memory, compared to 6.6 mins for svim and 0.34 GB memory. ONT reads at 13× coverage were analysing by dysgu in 59 mins using 0.94 GB memory, which was slower than the fastest caller svim (32 mins and 0.9 GB memory).

**Table 6.** Resource requirements of SV callers. Best scores are shaded blue.

| Reads | Caller | Mins | Mem (GB) |
| --- | --- | --- | --- |
| Illumina 40X | dysgu | 75.3 | 5.58 |
|  | manta | 365.1 | 0.24 |
|  | delly | 150.0 | 6.42 |
|  | lumpy | 211.5 | 12.00 |
| PacBio 8X | dysgu | 8.0 | 0.35 |
|  | nanovar | 46.5 | 19.05 |
|  | svim | 6.6 | 0.34 |
|  | sniffles | 20.3 | 0.71 |
| ONT 13X | dysgu | 59.7 | 0.94 |
|  | nanovar | 83.4 | 17.58 |
|  | svim | 32.4 | 0.90 |
|  | sniffles | 64.6 | 2.01 |

## Discussion

We developed dysgu to facilitate SV and indel discovery using PE or LR sequencing platforms in a computationally efficient manner. Dysgu analyses several forms of evidence to detect events including alignment gaps, discordant reads, read-depth, soft-clipped and supplementary mappings. For PE data, remapping of anomalous soft-clipped reads is also utilized to identify additional small SVs. Putative events are then labelled with a useful probability value using a gradient boosting classifier (28). Stratifying events by probability has several potential benefits over manually filtering. For example, machine learning classifiers can learn non-linear relationships between variables, and potentially capture large numbers of interactions between variables that would be difficult to reproduce through a manual approach. However, machine-learning raises additional challenges such as feature engineering, collation of appropriate training sets, and assessing how well a model will generalize to new data.

Dysgu models SV events using a vector of up to 41 features depending on read-type, with each feature designed to quantify different aspects of an SV signature, or error patterns of the respective read-type. The current list of features is non-exhaustive and can potentially be expanded in future releases to enhance calling performance.

Features incorporate more obvious signals such as read-support and sequencing depth, as well as novel patterns such as “soft-clip quality correlation” (PE data only) and repetitiveness scores (See Methods). To facilitate the calculation of features which capture sequence-contextual information, we also developed a novel linear-time consensus sequence algorithm, which is used to rapidly collapse reads at each break site into consensus contigs for further analysis. We trained our classifier using a large collection of manually labelled SV loci and combined these sites with loci identified by other SV callers. Manually labelling induces an obvious bias in the training set, where the correctness is a matter of opinion of the human observer. However, using a manual approach also allowed us to generate training sets with high completeness, which was not the case when relying on third party SV callers. Construction of quality training sets is a perennial challenge in machine learning and we expect that improving the quality and size of training sets will yield further performance improvements for SV classification.

We validated performance using benchmark datasets provided by GIAB (16, 17), and provide a software library ‘svbench’ to facilitate benchmarking and exploration of results. Primarily we assessed the HG002 benchmark, analysing in detail high-confidence Tier 1 regions, as well as all genomic regions. At Tier 1 regions we find that dysgu outperforms existing tools for both PE reads (Table 1) or third generation long-reads (Table 3,) using the F1 metric for comparisons. Tier 1 regions cover 2.51 Gbps of the genome although more complicated regions and smaller indel SVs (< 50 bp) are absent. Analysis of all genomic regions largely supported the conclusion that dysgu matches or outperforms existing tools, with dysgu often showing the best F1 scores across read types (Supplemental_Table_S6.pdf, Supplemental_Table_S7.pdf). Notably, svim showed higher F1 scores than dysgu in some benchmarks, although this was at the expense of considerably lower precision values and often increased duplication of true-positives.

Another novel feature of dysgu is that calls from separate sequencing technologies can be merged using a single command. Particularly, we found that adding calls made using Illumina data to either PacBio or ONT led to improved recall (Table 5). However, this appeared to occur mainly outside Tier 1 regions, suggesting Tier 1 regions are an ‘easy-case’ for LR platforms. Nevertheless, for applications that require higher recall, adding PE data to lower coverage LR data is a cost-effective approach for SV discovery that dysgu can support.

In conclusion, dysgu is de novo SV caller that outperforms existing tools using PE or LR datasets.

Dysgu is also computationally efficient to run, being the fasted tool using PE data, or second fastest using LR data. We provide dysgu as an open-source package for use in basic and applied research applications.

## Materials and methods

### Overview

Dysgu has been designed to work with aligned reads in BAM or CRAM formats, and can analyse PE reads with lengths in the range 100 – 250 bp, or single-end LR such as PacBio Sequel II, or ONT. By default, events with a minimum size of ≥ 30 bp are reported. Depending on the sequencing platform, dysgu offers pre-set options which apply recommended settings and a specific machine learning model (e.g. use ‘– mode pe’ or ‘—mode pacbio’ for PE or PacBio settings, respectively).

Dysgu provides a ‘run’ command which will produce a vcf file for a single input file, which is recommended for PE reads. However, depending on read-type the stages of the pipeline can differ. For PE reads (and optionally long reads), dysgu first partitions SV candidate reads into a temporary uncompressed bam file, which is achieved using the ‘fetch’ command. As this stage is time-consuming, this command can also be run in a stream during BAM file processing to further save wall runtime. Dysgu will then apply the ‘call’ command to SV candidate reads and produce an output. Depending on the length of input reads, the ‘fetch’ command may be redundant, as for very long reads such as ONT, a large proportion of reads harbour multiple SV candidates, which effectively leads to the input file being duplicated. Therefore the ‘fetch’ command is not needed for some LR datasets, and the ‘call’ command is recommended instead.

### Identifying SV candidate reads

For PE reads, library insert metrics are collected from the input file by scanning the first 200 x 10^3^ reads. If the ‘fetch’ command is utilized, single reads, or all alignments from a read-pair, that are deemed to be candidates, are partitioned into a temporary file. However, if the ‘fetch’ command is not run, then input reads are simply marked as SV candidates. A read is defined as a candidate if a read is found with either, map-quality ≥ 20, a soft-clip ≥ 15 bp (PE only), a discordant insert size or read orientation (PE only), a supplementary mapping, an alignment gap ≥ 30, or a mate on another chromosome. A discordant insert size is defined as *insert size ≥ insert median* + (5. *insert stdev*). Reads in high coverage regions of the genome are also not analysed by default, defined as regions with a mean depth ≥ 200 (‘--mode pe’) or ≥ 150 (‘—mode pacbio’ or ‘—mode nanopore’).

### Genome coverage

Dysgu collects several quality control metrics for use as features in the machine learning model. Genome coverage is calculated according to (29), except coverage is binned into 10 bp non-overlapping segments. The genome coverage tracks are saved in the temp folder during execution.

### Alignment clustering

Reads are initially clustered using an edge-coloured undirected graph *G*. Nodes in the graph represent SV-signatures and correspond to events listed in the cigar field of an alignment, or the properties of a read. SV-signatures are enumerated as either ‘discordant’, ‘split’, ‘deletion’, ‘insertion’ or ‘breakend’, and are associated with a ‘genomic-start’ and ‘genomic-end’ position. ‘Breakend’ types indicate a read that has a normal mapping orientation and no supplementary mappings, but has a soft-clipped sequence, which potentially corresponds to an unmapped breakpoint. Edges correspond to either ‘white edges’ that link together all alignments in a template with the same query name, or ‘black’ edges that are added between nodes that share a compatible SV signature.

Clustering is split into two phases. Initially, genomic reads are converted into a series of SV-signatures, with each item corresponding to a separate candidate event. For example, a deletion identified in the alignment cigar, a discordant read, or a read with an unmapped soft-clipped are converted into SV-signatures as nodes in *G*.

The local genomic region is then searched for events with a compatible signature. We use a red-black tree to search for items with a similar ‘genomic end’ position before checking if the ‘genomic start’ position is also similar. A search depth of 4 is used to search forwards and backwards in the data structure for other nodes. We find that using the ‘genomic end’ position permits a shallow search depth as datapoints are often sparser at the distant ‘genomic end’ position. Edges are not permitted between ‘deletion’ or ‘insertion’ types, although edges between other types are allowed.

When searching for other nodes to add ‘black’ edges between, nodes that are closer in the genome to the query are preferred, so if multiple candidates are found, edges are only formed between nodes passing a more stringent threshold. SV-signatures are checked to make sure that they have a reciprocal overlap of 0.1, and a separation distance between ‘genomic start’ and ‘genomic end’ positions below a clustering threshold. For PE reads, the clustering threshold is < *insert median* + (5. *insert stdev*) *bp*, while for PacBio the threshold is < 35 *bp*, and ONT < 100 *bp*. If another SV-signature is found with a ‘genomic start’ < 35 *bp*, these nodes pass the more stringent threshold, and a ‘black’ edge is added to the graph. For single-end reads or ‘split’ reads, if any of these conditions fail we also check the span position distance (26) between signatures. Span position distance between signatures S_1_ and S_2_ is defined as 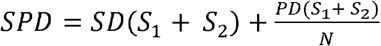 where SD is the span distance between signatures 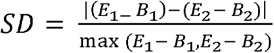, and PD is the position distance 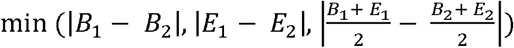. N is a normalization constant which is set at 100 for PE reads, 600 for PacBio and 900 for ONT reads. For all read types the SPD threshold used is *t* < 0.3. For PE reads that do not have a ‘split’ SV signature, we use a modified formula, only adding ‘black’ edges between nodes if 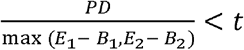 and *SD* < *t*.

If no edges are found for a PE read, a second phase of clustering is used to try and find edges between reads that share similar soft-clipped sequences. As pairwise sequence comparison between neighbouring alignments is computationally costly, we devised a novel algorithm based on clustering of the minimizer sketch of soft-clipped reads (30). Minimizer sampling involves computing the list of minimum kmers derived from consecutive windows over a sequence. We use a kmer length of 6 and a window length of 12. The minimum kmer is selected using a hash function and computed in linear-time O(n)(31). Additionally, in a modification of the minimizer sketching algorithm, we compute only the unique set of minimum kmers *S_k_* for each soft-clipped portion of a read. Each kmer in the set *S_k_* is associated with a genomic position that corresponds to the left-most or right-most base in the alignment for left or right soft-clipped sequences, respectively.

Kmers are added to a hashmap *M* with the key given by the kmer hash, and the value pair corresponding to a set of tuples, of (genomic position, read name). Kmers that are > 150 bp from the query genomic position are dynamically removed from the hashmap during processing.

For each incoming read, the kmer set *S_k_* is first computed, then for each kmer a corresponding set *Z* of reads and genomic positions is obtained by indexing *M*. The set *Z* consists of a collection of local reads that share the same minimizer kmer as the query. Entries in *Z* are then compared to the current genomic position and if the separation is < 7 bp, the number of found minimizers *a* is incremented. Additionally, the number of minimizers shared between reads with the same name *b* is counted. The total minimizer support is defined as 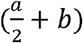 and a threshold of ≥ 2 is utilized. Once the minimizer support threshold is exceeded, found nodes are added to a set and returned.

Finally, ‘black’ edges are added to the graph between the returned set of nodes and the query node. Utilizing the minimizer clustering algorithm, pairwise sequence alignment is avoided, instead sequence matches between two sequences can be inferred from computing a minimizer sketch and utilizing hashmap queries.

### Event partitioning

Once all alignments have been added into the main graph *G*, the graph is simplified to a undirected quotient graph *Q* = (*V_q_,E_q_*) whose vertices consists of blocks or partitions of vertices from the main graph *G*. The vertices (partitions) *V_q_* are found by finding connected components in *G* using ‘black’ edges only. Edges *E_q_* are then defined between partitions using ‘white’ edge information from *G*, thus linking together read templates that map one or more SV.

Connected components in *Q* are processed together. These components can be composed of one or more partitions, harbouring potentially multiple SV events. In the simplest case, a component will consist of a single partition, which is processed for one or more SV. Components with a single edge are processed for a single SV only. For components with multiple edges, each edge is processed for a single SV, and additionally, each node partition is processed as a single partition if the number of ‘black’ intra-partition edges exceeds the number of ‘white’ out-edges, according to the main graph *G*. Thus, all components of *Q* are processed as a series of single-edges or single-partitions.

Single-edges in *Q* are assumed to represent a single SV, with reads from the *u* partition corresponding to one breaksite and reads from the *v* partition corresponding to the other. Single-partition nodes are assumed to map a single SV if a spanning alignment is found (e.g., a deletion event in the alignment cigar field). If no-spanning alignments are found, reads in the single-partition are further clustered using hierarchical clustering with the Nearest Point Algorithm (32), using the genomic start and end points of reads in the partition. This step is helps disentangle SVs with large overlaps and similar reference coordinates. Identified sub-clusters are then processed for a single SV.

### Consensus sequence generation

We generate consensus sequences at each breakpoint, from which read properties can be derived, such as repeat score or expanded polymer bases (see SV metrics section for further details), and to determine soft-clipped sequences for potentially remapping to the reference genome. We utilize a novel algorithm that borrows concepts from the positional de Brujin graph (33), and partial order alignment graphs (POA) (34). In a positional de Brujin graph *G*, the vertex set *V* encodes each sequence kmer in addition to genomic location, which helps leverage information provided by the mapper and localizes assembly. Edges *E* are permitted between kmers adjacent in the reference genome, which generally leads to a directed acyclic graph. However, it is possible that some bases do not have a genomic location, such as insertions within a read, or soft-clipped sequence. In such cases, genomic location can be inferred, for example using the expected mapping position if the whole read was aligned without gaps (10).

Partial order alignment graphs (34) are used to perform multiple sequence alignments, with vertices representing bases, and edges added between neighbouring bases in a sequence. Additional Sequences can be pairwise-aligned and incorporated into a POA using dynamic programming, and a consensus can be extracted by back-tracing through the maximum weighted path (34).

In our algorithm, we also represent vertices as bases and employ back-tracing through the longest path. However, similar to a positional de Brujin graph, we take the ordering of the graph from the genomic locations determined by the mapper. Utilizing this approach gives an approximation of a multiple sequence alignment between local genomic reads, and makes usage of information given by the mapper, whilst being simple and efficient to compute.

Let vertices correspond to a tuple (*b_i_,i,f,c*) ∈ *V*, where *b_i_* is the base aligned at genome position *i*, *i* is the genome position, *f* is an offset describing the distance to the closest aligned base, and *c* is a flag to indicate if the base is part of a left or right soft-clip (or neither). For left soft-clipped bases *c* = 1, right soft-clipped bases *c* = 2, whilst *c* = 0 otherwise. Bases that are not aligned to the reference genome may thus belong to three categories, when *f* > 0, for insertions *c* = 0, for left soft-clips *c* = 1, and for right soft-clips *c* = 2.

Edges are added between adjacent bases in a sequence (*u_j_*,*v*_*j*+1_), and vertices are weighted according to the sum of base qualities for a given node. Graph construction leads to a directed acyclic graph, that is then topologically sorted in linear time (35). To read the consensus sequence, the graph is first traversed using breadth-first search and for each vertex *v*, the longest path ending at *v* is determined by choosing the highest scoring predecessor vertex and adding to the running total. The consensus sequence is read by back-tracing from the vertex with the highest score, and recursively selecting the best predecessor node.

The worst-case time complexity for consensus sequence generation is linear with the number of input sequence bases. This follows, as graph construction, topological sorting, breadth-first search and back-tracing all have worst case complexities of *O*(*V* + *E*) time.

### Consensus sequence quality trimming

For the described consensus sequence algorithm, problems can arise at unmapped bases (e.g. soft-clipped sequences) if the underlying reads have a high indel error rate. In this situation, indels in unaligned bases cause neighbouring sequences to be shifted out of sync and can result in collapsing of indel errors in the consensus sequence. To address this problem, we trim soft-clipped sequences at bases with an alternative high scoring path. For each node *v* on the consensus path, with predecessor *u* and successor *w* also on the consensus path, a path quality metric is calculated. *l_total_* is defined as the total weight of all incoming edges to *v*. The in-edge quality is defined as 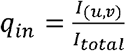, where *I_(u,v)_* is the weight of the consensus path edge (*u,v*). Similarly, *O_total_* is defined as the total weight of all outgoing edges from *v*. The out-edge quality is defined as 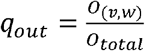, where *O_(v,w)_* is the weight of (*v,w*). The path quality metric for *v* is defined as *P_q_ = min(q_in_,q_out_)*. Soft-clipped sequences are trimmed at bases with a path quality metric < 0.5.

The soft clip weight (scw) parameter is defined for subsequent filtering, as the total base quality of nodes in the soft-clipped portion of the sequence divided by the length of the soft-clip.

### Re-mapping of contigs

After generating consensus sequences, if an end co-ordinate could not be determined, an attempt is made to align the soft-clipped sequence to the reference genome. Soft-clipped sequences are remapped to a window ±500 bp from the anchored breakpoint. We utilize edlib (36) (parameters: mode=“HW”) to find an approximate location, before refining the alignment using Striped Smith-Watermen (37) (parameters: match_score=2, mismatch_score=-8, gap_open_penalty=6, gap_extend_penalty=1) using the scikit-bio library (found online at: http://scikit-bio.org/). For deletion events, if less than 40 % of the soft-clip could be remapped and the alignment span is < 50bp, the alignment is rejected. For insertion events, if > 20 bp of sequence could not be mapped the alignment is rejected.

If no alignment is identified, dysgu can still call an unanchored insertion event at the identified break point, however, only events that have support > min_support + 4 and a soft-clip length ≥ 18 bp. The min_support parameter can be user supplied and takes a value of 3 for PE data or 2 for LR data.

### Sequence repeat score

Dysgu calculates repetitiveness scores for aligned regions of contigs as well as reference bases between deletions, and soft-clipped sequences. To calculate this metric, the sequence of interest is broken into kmers of increasing lengths from 2 - 6 bases. For each kmer of length *k*, a hashtable is used to record the last seen position of each kmer. If a kmer is seen more than once, the distance in bases to the last seen position is retrieved *d*. The repeat score is then calculated as a mean according to 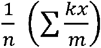 where *k* is the kmer length, and *x* and *m* have the form 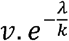, where *e* is Euler’s number, *λ* is a decay constant set at 0.25, and *v* = *k* for the denominator *m*, and *v* = *d* for *x*. For perfect tandem repeats 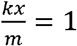, whilst sequencing errors, interspersed patterns or random sequence lead to lower values.

### Base quality score correlation at soft-clipped reads

For short-read input data we calculate a metric referred to as ‘soft-clip quality correlation’ (SQC), which is aimed at quantifying a sequence-specific error profile we observed in Illumina data (38). During sequencing, it is though that certain genomic sequences can promote dephasing, that gives rise to read base-qualities that correlate with the underlying sequence, and can result in frequent mismatches in alignments at specific bases (38). In our data, we observed a pattern consistent with this model but occurring at soft-clipped reads. These sites were frequently identified adjacent to homopolymer sequences and displayed base-quality scores that fluctuated with the underlying soft-clipped sequence. These soft-clip sequences often appeared to contain many errors as neighbouring soft-clipped reads showed many differences. Finally, these sites also frequently gave rise to false-positive calls at one-end anchored SV calls. The SQC metric was devised to quantify this phenomenon and is utilized as a feature in machine learning classification.

For each query read from the putative SV, the quality values of soft-clipped bases are added to a hashmap *H*, with the relative genomic position *pos* as the key, and a list *L_pos_* of base-qualities as values. The relative genomic position is taken as the position of the base if the whole soft-clipped portion of the read was mapped to the genome. Once all reads have been added, the ‘local mean’ is calculated as the absolute difference from the mean of each list *d_pos_* = |*x_j_* - *μ*| where *x_j_* is each item in *L_pos_* and *μ* is the mean of *L_pos_*. The sum of all calculated values of *d_pos_* is stored in a variable *v_local_* = *∑d_pos_*, and the global mean across all *d_pos_* is calculated 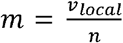. Finally, for each list in *H*, the sum of differences with the global mean is calculated *v_global_* = ∑|*x_j_* − *m*|. The SQC metric is calculated as the ratio 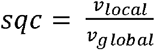. When the positions of low-quality bases are distributed randomly with genomic position *sqc* values will be close to 1.0. However, when low quality bases are clustered at certain positions, this results in smaller differences in base qualities at the local scale, giving smaller *v_local_* values and lower *sqc* values.

### Fold change in coverage across SVs

We calculate the fold change in coverage (FCC) across putative SVs according to (39) with minor modifications. We utilize a genomic bin size of 10 bp and analyse 1 kb sequence flanking the left and right breaksites. The fold change in coverage is calculated as the median coverage of the interior SV region divided by the median of the flanking sequence. The FCC metric was the most important feature after SV length for classifying SVs by machine learning, however we considered that this metric may not be suitable for non-diploid samples, or complex clonal mixtures such as those encountered during tumour sequencing, as lower allelic fractions only give rise to small changes in FCC. For this reason, we also provide an additional machine-learning model for use with non-diploid or complex tumour SV discovery.

### Polymer repeats at breaksites

Dysgu searches for simple repeat patterns with a unit length of 1-6 bp that directly overlap a break. These sites could arise from the joining of directed repeats (e.g. deletion event) or by the extension of the polymer at the break (e.g. insertion), or perhaps a more complex event. The length of the identified repeat sequence and the stride of the simple repeat are also utilized as features in the machine learning model.

For each base in the input sequence, a search is initiated for a repeat pattern starting at that base. Repeat lengths *l* of between 1-6 bp are tested in increasing length. To identify a repeat pattern, successive kmers are tested for identity with the starting kmer, using a step size of *l*. If a matching kmer is found the count *c* is incremented. If > 3 non-matching kmers or > 1 successive non-matching kmer is found the search is stopped. If *c* ≥ 3 when the search is stopped, and the spanning sequence identified is > 10 bp, the repeat sequence is set aside. Finally, if the repeat sequence overlaps the breaksite then the SV event is annotated with the breaksite repeat and stride length.

### SV event metrics

Dysgu annotates each putative SV event with a number of metrics. In Table 7, we list metrics utilized in the diploid paired-end model by decreasing feature importance.

**Table 7.**
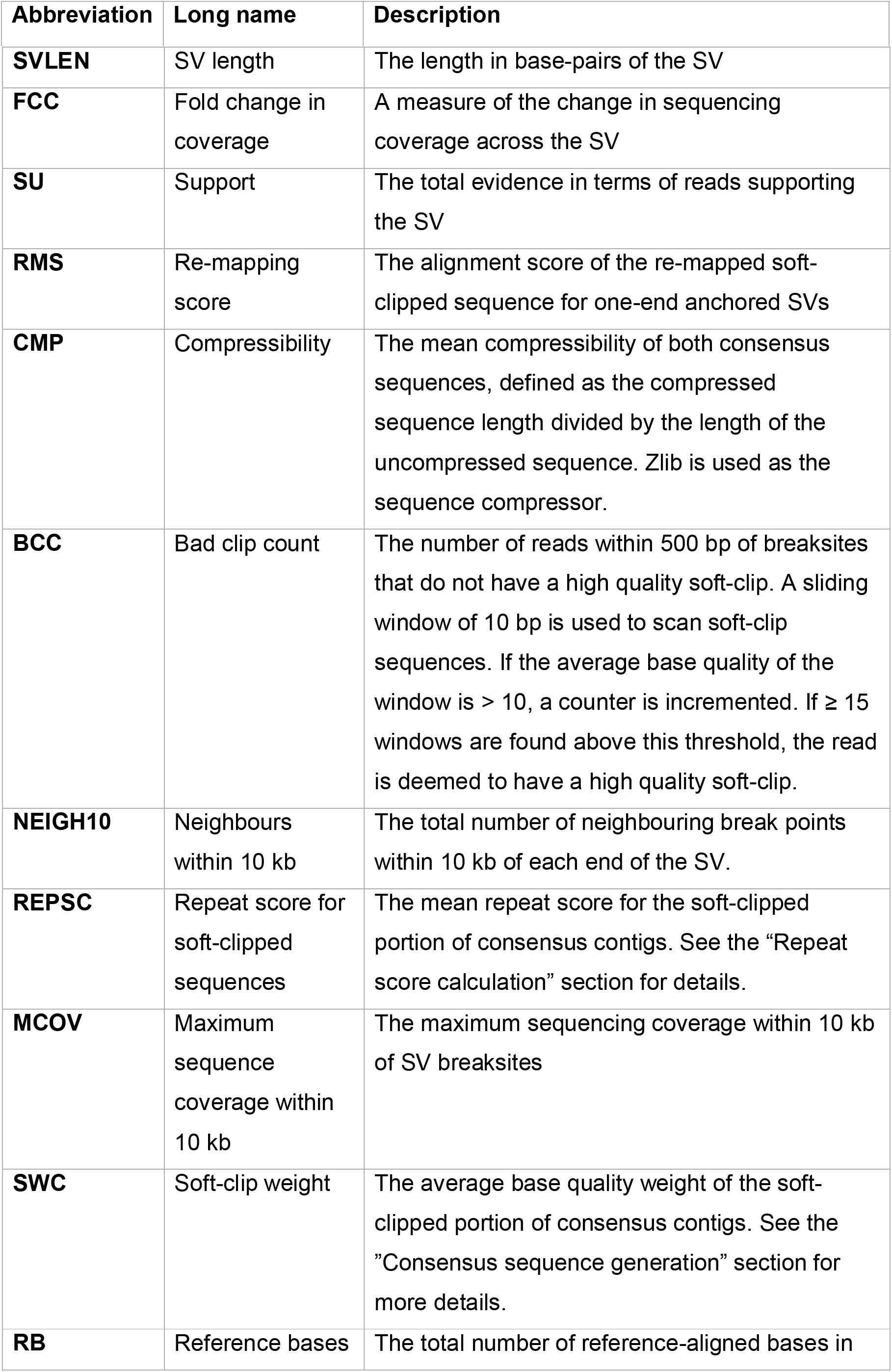
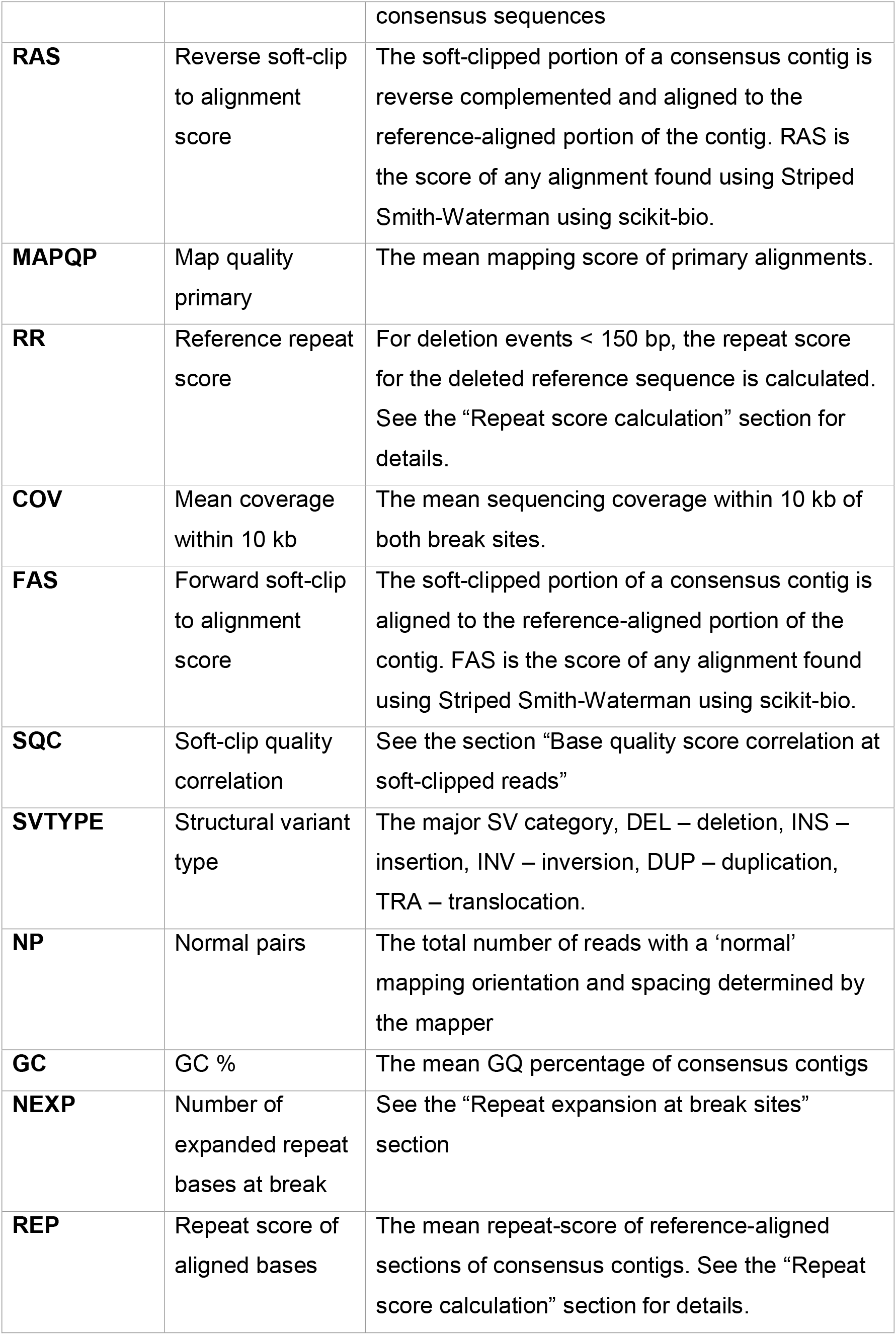
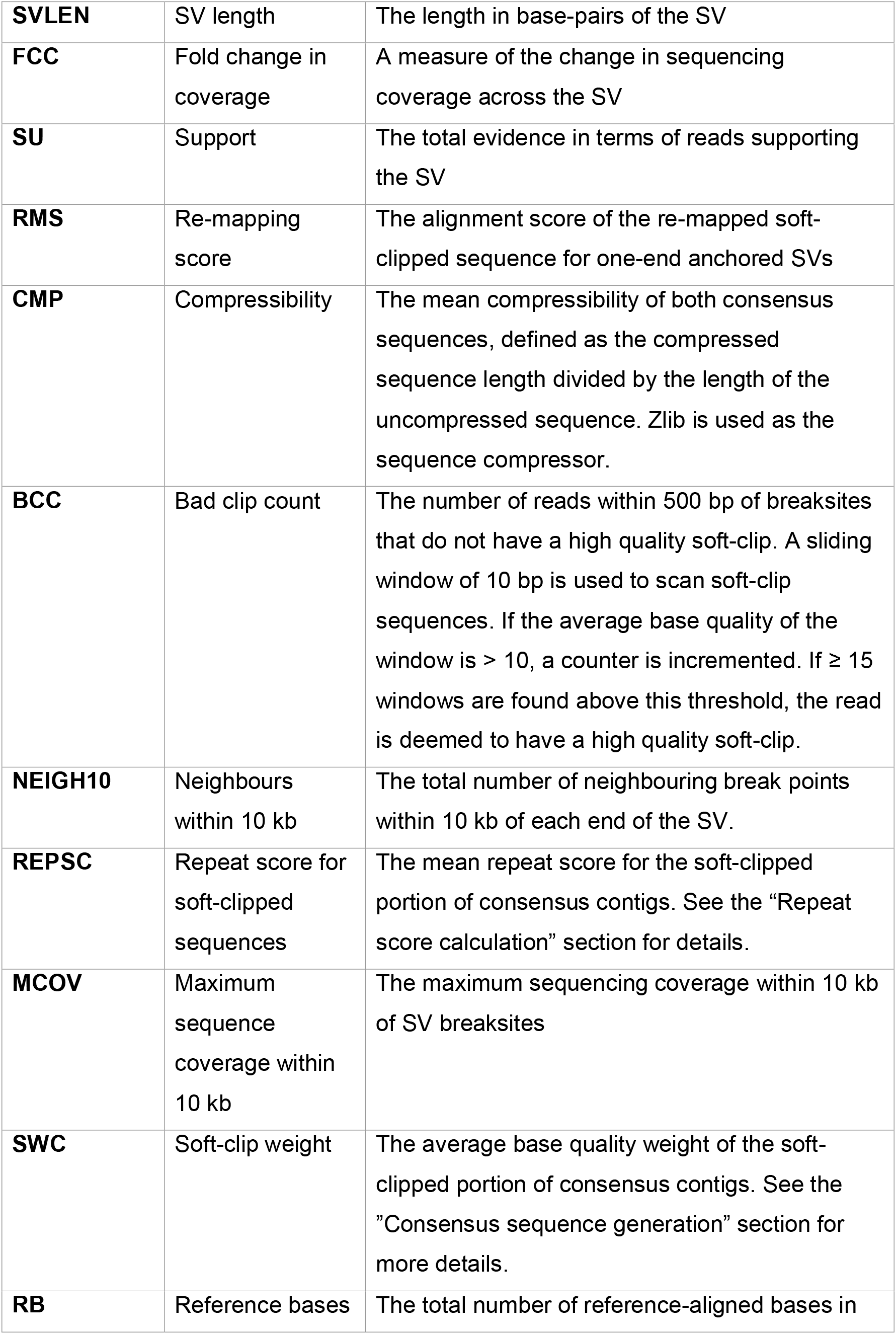
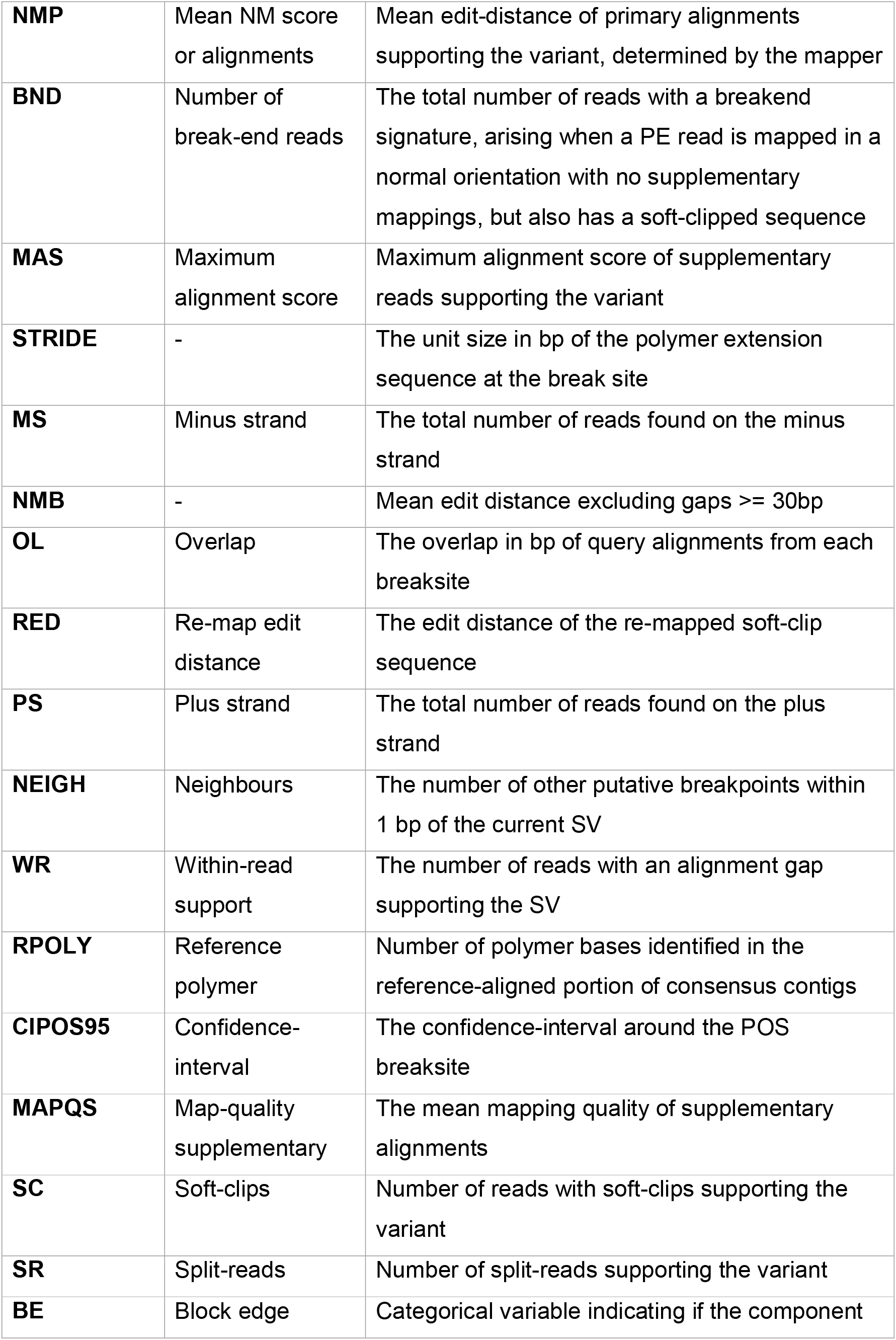
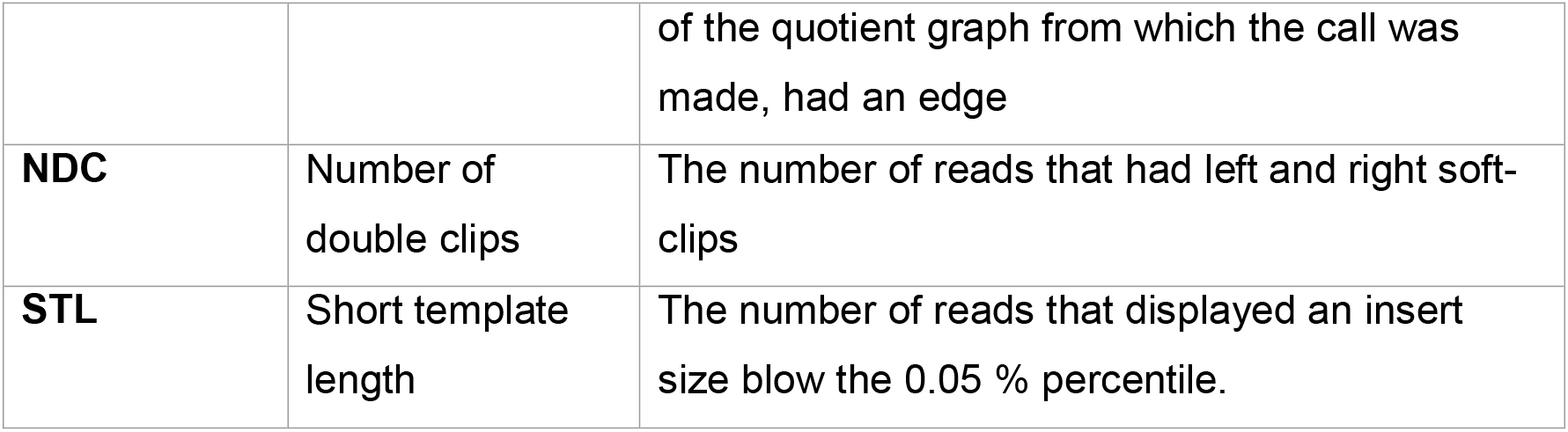
Overview of the features used in machine learning classification.

| Abbreviation | Long name | Description |
| --- | --- | --- |
| **SVLEN** | SV length | The length in base-pairs of the SV |
| **FCC** | Fold change in coverage | A measure of the change in sequencing coverage across the SV |
| **SU** | Support | The total evidence in terms of reads supporting the SV |
| **RMS** | Re-mapping score | The alignment score of the re-mapped soft-clipped sequence for one-end anchored SVs |
| **CMP** | Compressibility | The mean compressibility of both consensus sequences, defined as the compressed sequence length divided by the length of the uncompressed sequence. Zlib is used as the sequence compressor. |
| **BCC** | Bad clip count | The number of reads within 500 bp of breaksites that do not have a high quality soft-clip. A sliding window of 10 bp is used to scan soft-clip sequences. If the average base quality of the window is > 10, a counter is incremented. If $\geq 15$ windows are found above this threshold, the read is deemed to have a high quality soft-clip. |
| **NEIGH10** | Neighbours within 10 kb | The total number of neighbouring break points within 10 kb of each end of the SV. |
| **REPSC** | Repeat score for soft-clipped sequences | The mean repeat score for the soft-clipped portion of consensus contigs. See the "Repeat score calculation" section for details. |
| **MCOV** | Maximum sequence coverage within 10 kb | The maximum sequencing coverage within 10 kb of SV breaksites |
| **SWC** | Soft-clip weight | The average base quality weight of the soft-clipped portion of consensus contigs. See the "Consensus sequence generation" section for more details. |
| **RB** | Reference bases | The total number of reference-aligned bases in |
|  |  | consensus sequences |
| **RAS** | Reverse soft-clip to alignment score | The soft-clipped portion of a consensus contig is reverse complemented and aligned to the reference-aligned portion of the contig. RAS is the score of any alignment found using Striped Smith-Waterman using scikit-bio. |
| **MAPQP** | Map quality primary | The mean mapping score of primary alignments. |
| **RR** | Reference repeat score | For deletion events < 150 bp, the repeat score for the deleted reference sequence is calculated. See the “Repeat score calculation” section for details. |
| **COV** | Mean coverage within 10 kb | The mean sequencing coverage within 10 kb of both break sites. |
| **FAS** | Forward soft-clip to alignment score | The soft-clipped portion of a consensus contig is aligned to the reference-aligned portion of the contig. FAS is the score of any alignment found using Striped Smith-Waterman using scikit-bio. |
| **SQC** | Soft-clip quality correlation | See the section “Base quality score correlation at soft-clipped reads” |
| **SVTYPE** | Structural variant type | The major SV category, DEL – deletion, INS – insertion, INV – inversion, DUP – duplication, TRA – translocation. |
| **NP** | Normal pairs | The total number of reads with a ‘normal’ mapping orientation and spacing determined by the mapper |
| **GC** | GC % | The mean GQ percentage of consensus contigs |
| **NEXP** | Number of expanded repeat bases at break | See the “Repeat expansion at break sites” section |
| **REP** | Repeat score of aligned bases | The mean repeat-score of reference-aligned sections of consensus contigs. See the “Repeat score calculation” section for details. |
| **NMP** | Mean NM score or alignments | Mean edit-distance of primary alignments supporting the variant, determined by the mapper |
| **BND** | Number of break-end reads | The total number of reads with a breakend signature, arising when a PE read is mapped in a normal orientation with no supplementary mappings, but also has a soft-clipped sequence |
| **MAS** | Maximum alignment score | Maximum alignment score of supplementary reads supporting the variant |
| **STRIDE** | - | The unit size in bp of the polymer extension sequence at the break site |
| **MS** | Minus strand | The total number of reads found on the minus strand |
| **NMB** | - | Mean edit distance excluding gaps $\geq 30$bp |
| **OL** | Overlap | The overlap in bp of query alignments from each breaksite |
| **RED** | Re-map edit distance | The edit distance of the re-mapped soft-clip sequence |
| **PS** | Plus strand | The total number of reads found on the plus strand |
| **NEIGH** | Neighbours | The number of other putative breakpoints within 1 bp of the current SV |
| **WR** | Within-read support | The number of reads with an alignment gap supporting the SV |
| **RPOLY** | Reference polymer | Number of polymer bases identified in the reference-aligned portion of consensus contigs |
| **CIPOS95** | Confidence-interval | The confidence-interval around the POS breaksite |
| **MAPQS** | Map-quality supplementary | The mean mapping quality of supplementary alignments |
| **SC** | Soft-clips | Number of reads with soft-clips supporting the variant |
| **SR** | Split-reads | Number of split-reads supporting the variant |
| **BE** | Block edge | Categorical variable indicating if the component |
|  |  | of the quotient graph from which the call was made, had an edge |
| **NDC** | Number of double clips | The number of reads that had left and right soft-clips |
| **STL** | Short template length | The number of reads that displayed an insert size blow the 0.05 % percentile. |

### Classifier training

To train a machine learning classifier for the different read-types (PE, PacBio and ONT) we constructed several ‘gold-sets’. Gold-sets consisted of manually curated SV loci or SV loci found using other calling software. Primarily, gold-sets were based on the well-studied HG001 sample (Female, Western European ancestry). However, for PacBio data, gold-sets were also derived from the HG005 sample (Male, Chinese ancestry). The read data utilized in constructing the gold-sets are listed in Table 8.

**Table 8.**
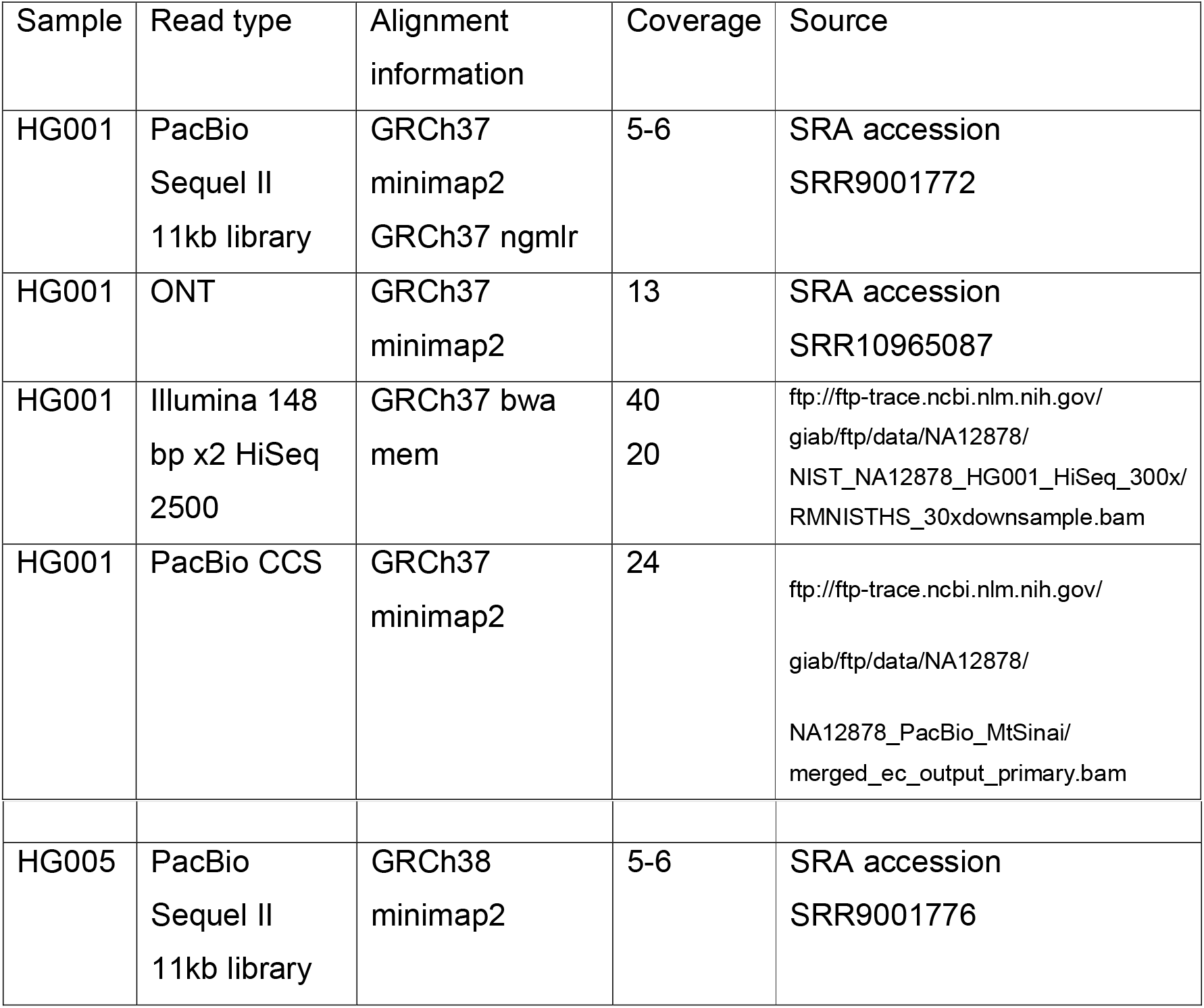
Overview of datasets used in model training.

The overall strategy was to quantify dysgu performance on smaller subsets of data, and then combine these smaller benchmarks into a larger set for training. We employed this strategy as it meant that manual curation of smaller subsets was more feasible (as opposed to annotating events genome wide), and also multiple methods for annotating true-positive calls could be integrated into the training set e.g. relying on manual curation, labelling using a third party SV caller, or utilizing previously publish call sets, or utilizing different DNA mappers.

Firstly, we constructed a gold-set based on PacBio Sequel II reads. Nanovar was run on HG001 minimap2-aligned reads and insertion calls from chr1 and chr10 in the size range 30-500 bp were added to the set (n=1808). The choice of chromosome to utilize was arbitrary. We also utilized a previously published list of deletion and insertion calls made using pbsv (n=27662) on PacBio CCS data at around 30× coverage (downloaded from GIAB ftp://ftp-trace.ncbi.nlm.nih.gov/giab/ftp/data/ChineseTrio/analysis/PacBio_pbsv_05212019/HG005_GRCh38.pbsv.vcf.gz).

Next we added a collection of manually curated SV loci that were identified by visually inspecting calls made by dysgu using the Integrative Genomics Viewer (IGV) (40). Multiple read-types were assessed, simultaneously viewing alignments of PacBio Sequel II, PacBio CCS and ONT reads. If the SV showed support in more than one technology the SV loci was labelled as true. If a call made by dysgu was plausible, but showed strong evidence of being below the minimum size threshold < 30 bp, then the call was labelled as false. All deletion and insertion calls for chr1, 10 and 11 for HG001 minimap2-aligned reads were manually labelled in this way (n=2973). Additionally, large insertion calls (“large-INS”) made by dysgu (≥ 500 bp, whole genome) using HG001 minimap2 and ngmlr aligned reads were also assessed (n=1661). Calls made by dysgu were then compared to these smaller benchmark sets separately and labelled as true or false using SVBench (available online at https://github.com/kcleal/svbench).

These smaller benchmarks were then concatenated before training a gradient boosting classifier using the lightgbm package (28) (boosting type “dart”). Features were first selected using recursive feature-selection with cross-validation using scikit-learn (41). Hyperparameters were tuned using grid search with cross-validation using Stratified K-fold (n=5) (41). The learning-rate, max-bin, max-depth, n-estimators and number-of-leaves were optimized in this way, whilst other parameters were left as default.

Events labelled using the PacBio classifier with probability ≥ 0.5 were then leveraged to help construct additional gold-sets for PE and ONT read-types. For the PE gold-set, deletion and insertion loci identified using the PacBio model were taken as true-positive loci (chromosomes 1, 2, 10, 11, 12, n=8258). Additionally, the “large-INS” set derived from PacBio reads was utilized. Finally, events called by dysgu using PE reads (HG001, bwa mem) were manually curated, corresponding to deletions (n=5984 true) from chromosomes 1 – 5 and 10 – 22, plus insertions (n=2250 true) from chromosomes 1-14. The choices of chromosomes were arbitrary.

For the ONT gold-set, we utilized deletion and insertion loci identified using the PacBio model (probability ≥ 0.5, whole genome n=25072 true). To this we used regions identified by Nanovar (n=23581 true), and the “large-INS” manually curated set. Additionally, we added manually curated dysgu calls from ONT data from chr1 and chr10 (n=4265).

### Benchmark datasets

For the HG002 benchmark, variants were downloaded from GIAB ftp://ftptrace.ncbi.nlm.nih.gov/giab/ftp/data/AshkenazimTrio/analysis/NIST_SVs_Inte_gration_v0.6. For HG001, variants were downloaded from GIAB ftp://ftp-trace.ncbi.nlm.nih.gov/giab/ftp/technical/svclassify_Manuscript/Supplementary_Information/Personalis_1000_Genomes_deduplicated_deletions.bed.

### Benchmarking SV calls using svbench

We developed a python software library “svbench” to facilitate rapid benchmarking of SV datasets, as well as to facilitate exploration and comparison of SV calls as an aide during software development. Svbench performs a similar role to other benchmarking programs such as truvari from GIAB (16), although as data structures can be held in memory and explored interactively, significant speedups can be obtained for benchmarking which can be helpful during software development and analysis.

Svbench also optionally adds a weighting to input SVs that can be used to break ties between multiple query and reference SVs. The weighting or “strata” can be specified during loading of SVs, and usually takes the value of a quality metric set by the caller, or if this is absent, the variant support in terms of read evidence. Stratifying SV calls in this way is also necessary to generate a precision-recall curve. Another difference between svbench and truvari, is that svbench can optionally classify duplicate true-positive calls, which can arise when one reference SV in the sample gives rise to multiple calls in the output. There are several ways to classify duplicates, such as labelling all duplicates as false-positives, true-positives, or ignoring them from precision calculation. By default, svbench utilizes the latter option. Although this can lead to optimistic precision and F1 scores, we consider this approach often leads to a clearer understanding of the underlying performance of an SV caller. For example, if duplicates are labelled as false-positives then a caller that identifies the correct genomic loci but has a high duplication rate is penalized, while a caller that identified incorrect loci but also has a low duplication rate could end up with a similar overall precision and F1 score. Furthermore, removing duplicates bioinformatically, might be less of a challenge than removing genuine false positives, by for example filtering SVs with low weight but found nearby other SVs.

Conceptually, svbench loads input files (vcf, bed, bedpe or csv format) into a ‘CallSet’ object. Internally, SV records are held in a pandas dataframe (42), which support a rich set of data wrangling capabilities, making common data operations straightforward such as filtering, splitting, combining, grouping, and plotting precision-recall curves.

To compare one dataset with another i.e. a benchmark dataset with a query dataset, both sets of SV loci are loaded into an svbench CallSet object. The benchmark dataset is then prepared by adding intervals (add_intervals function) around each breaksite, adding one interval for each start and end coordinate. Intervals are held in a nested containment list using the ncls library (43). Utilizing an interval at both start and end sites, rather than a single interval, means translocations can be naturally compared, and for large SVs, nesting of small SV intervals within larger SVs is avoided which can reduce the search space when comparing records.

Query SVs are then checked against prepared intervals. If a benchmark record overlaps both the start and end of a query SV, and the percent size similarity, reciprocal overlap and svtype match criteria, then the records are considered to match. Percent size is defined as 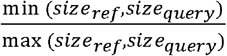. Query and benchmark records that pass provided thresholds are then clustered on an undirected graph *G*, using the network library (44).

Edges (*u,v*) ∈ *G* are added to the graph between benchmark vertices *u* and query vertices *v* with the edge weight given by the “strata”, or weight property of the query event, which is parsed during loading of the data. If a query vertex *v* matches multiple benchmark vertices *u*, then the chosen benchmark call *u* is determined by the closest absolute genomic distance between *u* and *v*, defined as |*start_query_* - *start_ref_*| + |*end_query_ - end_ref_*|. Once all query records have been added to the graph, connected components are then processed. If a benchmark vertex has multiple edges, a highest scoring edge is selected as the true-positive call, whilst other query vertices are labelled as duplicates. If duplicate classification is permitted then precision scores are calculated as 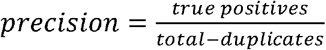. If duplicate classification is turned off then duplicates are treated as false positives. Recall is assessed as 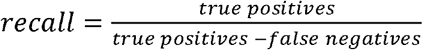 and F1 score is calculated as 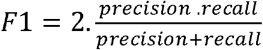.

We utilized svbench to assess performance of dysgu compared to other SV callers. For benchmarking calls against the HG002 benchmark (16), we filtered query calls by a minimum size of 30 bp (whole genome benchmark), or 50 bp (Tier 1 benchmark). We utilized a reference interval size of 1000 bp, and a percent size similarity threshold of 15 %. Deletion and insertion calls were analysed separately, filtering both query and reference calls by svtype before comparison. Additionally, only query calls on the ‘normal’ chromosomes were analysed {*chr*1..*chrY*}. To match the definition of the GIAB benchmark, we converted DUP calls < 500 bp to insertions.

SV callers were applied to datasets using default settings. Version numbers for tested callers were as follows: dysgu v1.1.4, gatk v4.1.2.0, strelka v2.9.2, manta v1.6.0, svim v1.3.1, sniffles v1.0.12, nanovar v1.3.2, delly v0.8.5. SV calls were also filtered by removing calls without a ‘PASS’ in the filter field (if applicable). The ‘strata’ metric utilized for each of the SV callers was as follows: lumpy – “SU”, delly – “QUAL”, dysgu – “PROB”, manta – “QUAL”, strelka – “QUAL”, gatk – “QUAL”, nanovar – “QUAL”, sniffles – “RE”, svim – “SUPPORT”. Events with a minimum support < 2 were filtered out.

## Abbreviations

SV: structural variant
PE: paired-end
LR: long-read
DEL: deletion
DUP: duplication
INV: inversion
INS: insertion
TRA: translocation
ONT: Oxford Nanopore Technologies
GIAB: Genome In A Bottle consortium
SRA: Sequencing read Archive
POA: partial order alignment

## Data availability

Dysgu is released as free and open source under the Massachusetts Institute of Technology (MIT) licence. Source code and distributions can be downloaded at https://github.com/kcleal/dysgu. Data used to train the classifier is available online at https://zenodo.org/record/4761527. Svbench is also released under the MIT license and can be found at https://github.com/kcleal/svbench. Analysis scripts used to reproduce results found in this paper can be found under https://github.com/kcleal/svbench. Illumina sequencing data for Ashkenazim HG002 (16) sample was downloaded from GIAB (ftp://ftp-trace.ncbi.nlm.nih.gov/giab/ftp/data/AshkenazimTrio/HG002_NA24385_son/NIST_HiSeq_HG002_Homogeneity-10953946/HG002Run01-11419412/HG002run1_S1.bam). Two lanes of PacBio data were downloaded from SRA (https://www.ncbi.nlm.nih.gov/sra) under accessions SRR10188368 and SRR10188369. ONT data were downloaded from SRA under accession SRR11537600.

## Author contributions

KC devised methodology, performed experiments, wrote the software and drafted the manuscript. DMB contributed design ideas, provided feedback and performed manuscript editing. All authors read and approved the final manuscript.

