## Supplemental_Fig_S1 for "Dysgu: efficient structural variant calling using short or long reads"

**Supplementary figures and tables**

### Illumina paired-end reads

|  | TP | | FP | | Precision | | Recall | | Duplication | | F1 | |
| --- | --- | --- | --- | --- | --- | --- | --- | --- | --- | --- | --- | --- |
|  | DEL | INS | DEL | INS | DEL | INS | DEL | INS | DEL | INS | DEL | INS |
| dysgu | 6955 | 4887 | 256 | 267 | 0.965 | 0.948 | 0.186 | 0.134 | 0.002 | 0.012 | 0.312 | 0.234 |
| manta | 3495 | 1364 | 169 | 23 | 0.954 | 0.983 | 0.094 | 0.037 | 0.002 | 0.011 | 0.170 | 0.072 |
| gatk | 4391 | 3225 | 215 | 280 | 0.953 | 0.920 | 0.117 | 0.088 | 0.022 | 0.011 | 0.209 | 0.161 |
| strelka | 2275 | 1977 | 44 | 233 | 0.981 | 0.895 | 0.061 | 0.054 | 0.002 | 0.019 | 0.115 | 0.102 |
| delly | 5209 | 537 | 980 | 12 | 0.842 | 0.978 | 0.139 | 0.015 | 0.003 | 0.000 | 0.239 | 0.029 |
| lumpy | 2793 |  | 939 |  | 0.748 |  | 0.075 |  | 0.003 |  | 0.136 |  |

Table S1. 20X Illumina aligned with bwa mem, All-regions benchmark.

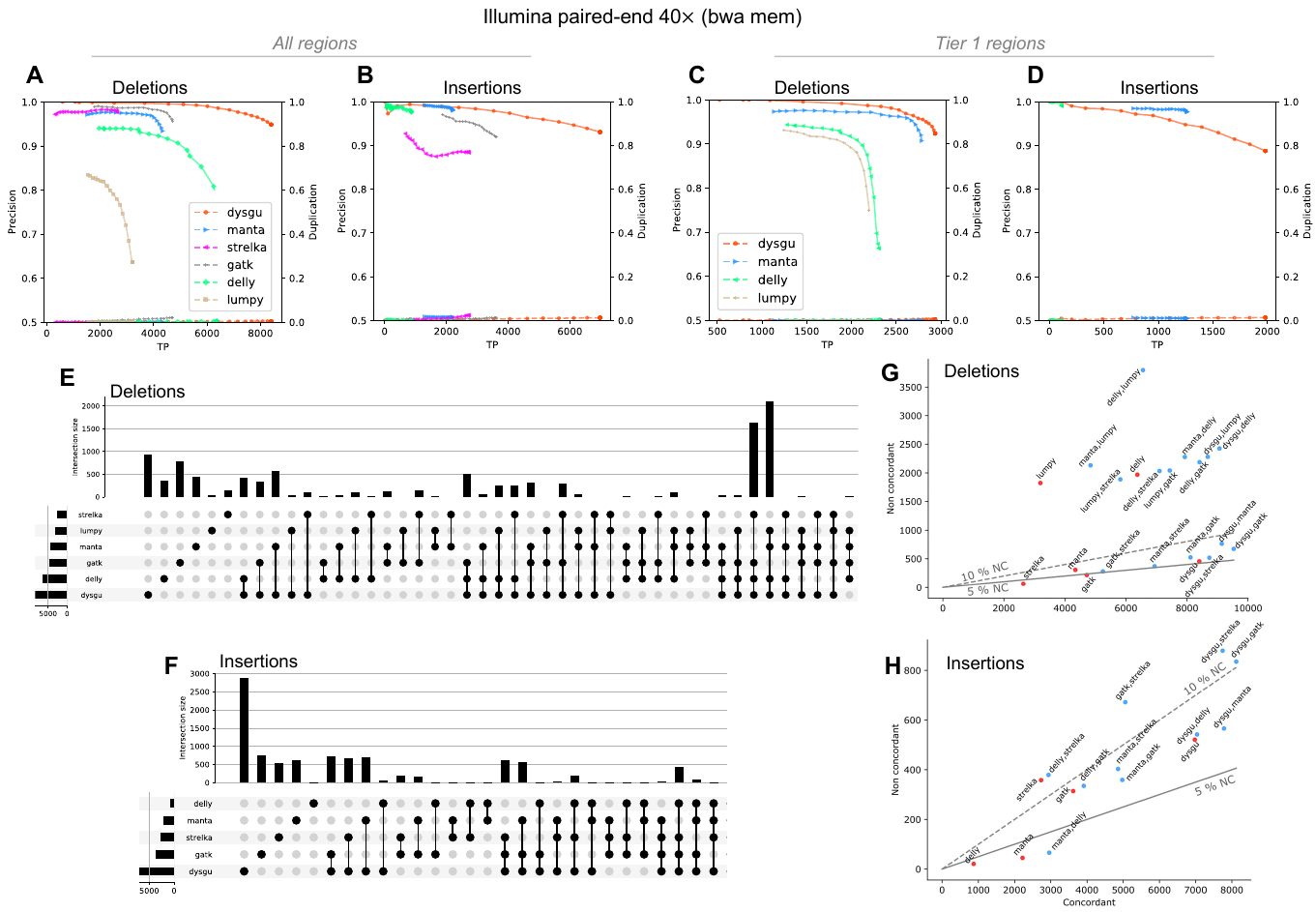

****Figure S1. Performance of dysgu using 40****$\boldsymbol{\times}$****PE reads.****

Precision-recall curves are illustrated for all genomic regions (A, B), and high-confidence Tier 1 regions (C, D). A secondary y-axis indicates duplicate true-positives (TP) as a fraction of true-positive calls. Intersections and aggregates of intersections of SV calls for the all-regions benchmark are shown as an upset plot (E, F). Combinations of callers are shown by plotting the union of true-positives between callers (labelled concordant), against the sum of false-positives (labelled non concordant) (G, H). The 5 and 10 % non-concordance (NC) is shown as a solid or dashed line, respectively.

|  | TP | | FP | | Precision | | Recall | | Duplication | | F1 | |
| --- | --- | --- | --- | --- | --- | --- | --- | --- | --- | --- | --- | --- |
|  | DEL | INS | DEL | INS | DEL | INS | DEL | INS | DEL | INS | DEL | INS |
| dysgu | 2936 | 1982 | 242 | 251 | 0.924 | 0.888 | 0.696 | 0.364 | 0.004 | 0.013 | 0.794 | 0.516 |
| manta | 2786 | 1265 | 285 | 30 | 0.907 | 0.977 | 0.661 | 0.232 | 0.000 | 0.009 | 0.765 | 0.376 |
| delly | 2305 | 112 | 1172 | 1 | 0.663 | 0.991 | 0.547 | 0.021 | 0.003 | 0.000 | 0.599 | 0.040 |
| lumpy | 2199 |  | 734 |  | 0.750 |  | 0.522 |  | 0.002 |  | 0.615 |  |

Table S2. 40X Illumina aligned with bwa mem, Tier1 benchmark.

|  | TP | | FP | | Precision | | Recall | | Duplication | | F1 | |
| --- | --- | --- | --- | --- | --- | --- | --- | --- | --- | --- | --- | --- |
|  | DEL | INS | DEL | INS | DEL | INS | DEL | INS | DEL | INS | DEL | INS |
| dysgu | 8587 | 6864 | 473 | 484 | 0.948 | 0.934 | 0.230 | 0.188 | 0.004 | 0.013 | 0.370 | 0.313 |
| manta | 4363 | 2220 | 280 | 42 | 0.940 | 0.981 | 0.117 | 0.061 | 0.003 | 0.014 | 0.208 | 0.114 |
| gatk | 4727 | 3638 | 207 | 296 | 0.958 | 0.925 | 0.126 | 0.100 | 0.022 | 0.014 | 0.223 | 0.180 |
| strelka | 2642 | 2747 | 62 | 342 | 0.977 | 0.889 | 0.071 | 0.075 | 0.002 | 0.024 | 0.132 | 0.139 |
| delly | 6460 | 872 | 1876 | 19 | 0.775 | 0.979 | 0.173 | 0.024 | 0.006 | 0.000 | 0.283 | 0.047 |
| lumpy | 3267 |  | 1749 |  | 0.651 |  | 0.087 |  | 0.005 |  | 0.154 |  |

Table S3. 40X Illumina aligned with bwa mem, All-regions benchmark.

|  |  | Precision | | | | Recall | | | | F1 | | | |
| --- | --- | --- | --- | --- | --- | --- | --- | --- | --- | --- | --- | --- | --- |
|  |  | [30, 50) | [50, 500) | [500, 5000) | ≥5000 | [30, 50) | [50, 500) | [500, 5000) | ≥5000 | [30, 50) | [50, 500) | [500, 5000) | ≥5000 |
| Deletions | dysgu | 0.94 | 0.94 | 0.99 | 0.94 | 0.45 | 0.29 | 0.38 | 0.34 | 0.61 | 0.44 | 0.55 | 0.5 |
|  | manta | 1 | 0.94 | 0.95 | 0.79 | 0.01 | 0.28 | 0.33 | 0.36 | 0.03 | 0.43 | 0.49 | 0.49 |
|  | gatk | 0.97 | 0.94 | 1 |  | 0.39 | 0.11 | 0 |  | 0.55 | 0.2 | 0 |  |
|  | strelka | 0.98 | 1 |  |  | 0.3 | 0 |  |  | 0.46 | 0.01 |  |  |
|  | delly | 0.95 | 0.77 | 0.64 | 0.26 | 0.34 | 0.19 | 0.41 | 0.42 | 0.5 | 0.3 | 0.5 | 0.32 |
|  | lumpy | 0.69 | 0.84 | 0.62 | 0.21 | 0 | 0.18 | 0.41 | 0.43 | 0.01 | 0.29 | 0.5 | 0.28 |
| Insertions | dysgu | 0.95 | 0.89 | 0.98 | 1 | 0.39 | 0.21 | 0.13 | 0.11 | 0.55 | 0.34 | 0.23 | 0.19 |
|  | manta | 0.98 | 0.98 | 1 | 1 | 0.02 | 0.16 | 0.01 | 0 | 0.03 | 0.28 | 0.03 | 0.01 |
|  | gatk | 0.94 | 0.89 | 1 | 1 | 0.27 | 0.12 | 0.02 | 0.03 | 0.42 | 0.21 | 0.03 | 0.05 |
|  | strelka | 0.87 | 0.93 | 1 |  | 0.31 | 0.01 | 0 |  | 0.46 | 0.02 | 0.01 |  |
|  | delly | 0.97 | 0.99 |  |  | 0.09 | 0.01 |  |  | 0.16 | 0.02 |  |  |

Table S4. 40X Illumina aligned with bwa mem, All-regions benchmark, split by size.

### PacBio Sequel II reads

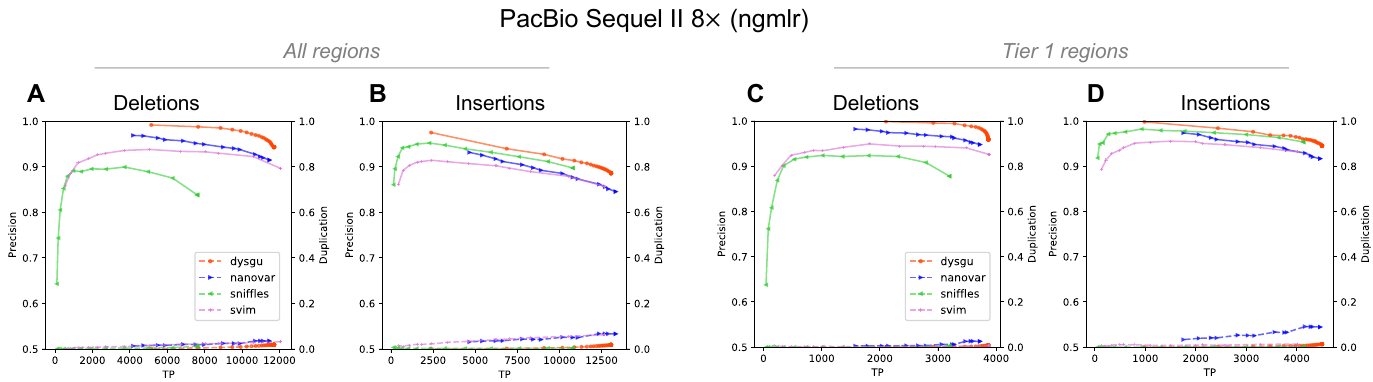

****Figure S2. Performance of dysgu on PacBio reads aligned using ngmlr.****

Precision-recall curves are shown for the ‘all-regions’ benchmark (A, B), as well as Tier 1 high-confidence regions (C, D).

|  | TP | | FP | | Precision | | Recall | | Duplication | | F1 | |
| --- | --- | --- | --- | --- | --- | --- | --- | --- | --- | --- | --- | --- |
|  | DEL | INS | DEL | INS | DEL | INS | DEL | INS | DEL | INS | DEL | INS |
| dysgu | 3894 | 4624 | 175 | 232 | 0.957 | 0.952 | 0.923 | 0.850 | 0.013 | 0.024 | 0.940 | 0.898 |
| nanovar | 3740 | 4502 | 169 | 332 | 0.957 | 0.931 | 0.887 | 0.827 | 0.030 | 0.096 | 0.921 | 0.876 |
| svim | 3893 | 4039 | 273 | 259 | 0.935 | 0.940 | 0.923 | 0.742 | 0.011 | 0.017 | 0.929 | 0.829 |
| sniffles | 3257 | 4146 | 367 | 182 | 0.899 | 0.958 | 0.772 | 0.762 | 0.011 | 0.002 | 0.831 | 0.849 |

Table S5. 8X PacBio Sequel II reads aligned using ngmlr2, Tier1 benchmark.

|  | TP | | FP | | Precision | | Recall | | Duplication | | F1 | |
| --- | --- | --- | --- | --- | --- | --- | --- | --- | --- | --- | --- | --- |
|  | DEL | INS | DEL | INS | DEL | INS | DEL | INS | DEL | INS | DEL | INS |
| dysgu | 12158 | 14212 | 864 | 2104 | 0.934 | 0.871 | 0.325 | 0.389 | 0.035 | 0.041 | 0.482 | 0.537 |
| nanovar | 11643 | 13834 | 872 | 1976 | 0.930 | 0.875 | 0.311 | 0.378 | 0.038 | 0.076 | 0.467 | 0.528 |
| svim | 12493 | 14718 | 2730 | 3814 | 0.821 | 0.794 | 0.334 | 0.403 | 0.067 | 0.170 | 0.475 | 0.534 |
| sniffles | 7929 | 12309 | 1581 | 1807 | 0.834 | 0.872 | 0.212 | 0.337 | 0.026 | 0.018 | 0.338 | 0.486 |

Table S6. 8X PacBio Sequel II aligned using minimap2, All-regions benchmark.

|  | TP | | FP | | Precision | | Recall | | Duplication | | F1 | |
| --- | --- | --- | --- | --- | --- | --- | --- | --- | --- | --- | --- | --- |
|  | DEL | INS | DEL | INS | DEL | INS | DEL | INS | DEL | INS | DEL | INS |
| dysgu | 11765 | 13190 | 504 | 1418 | 0.959 | 0.903 | 0.315 | 0.361 | 0.022 | 0.026 | 0.474 | 0.516 |
| nanovar | 11594 | 13478 | 884 | 2154 | 0.929 | 0.862 | 0.310 | 0.369 | 0.040 | 0.076 | 0.465 | 0.516 |
| svim | 12184 | 12814 | 1151 | 1885 | 0.914 | 0.872 | 0.326 | 0.350 | 0.040 | 0.072 | 0.480 | 0.500 |
| sniffles | 7899 | 10880 | 1096 | 1114 | 0.878 | 0.907 | 0.211 | 0.298 | 0.022 | 0.006 | 0.341 | 0.448 |

Table S7. 8X PacBio Sequel II aligned using ngmlr, All-regions benchmark.

|  |  | Precision | | | | Recall | | | | F1 | | | |
| --- | --- | --- | --- | --- | --- | --- | --- | --- | --- | --- | --- | --- | --- |
|  |  | [30, 50) | [50, 500) | [500, 5000) | ≥5000 | [30, 50) | [50, 500) | [500, 5000) | ≥5000 | [30, 50) | [50, 500) | [500, 5000) | ≥5000 |
| Deletions | dysgu | 0.951 | 0.964 | 0.962 | 0.844 | 0.515 | 0.482 | 0.453 | 0.354 | 0.668 | 0.642 | 0.616 | 0.499 |
|  | nanovar | 0.937 | 0.943 | 0.904 | 0.521 | 0.487 | 0.470 | 0.452 | 0.374 | 0.641 | 0.627 | 0.603 | 0.436 |
|  | svim | 0.915 | 0.924 | 0.864 | 0.672 | 0.530 | 0.498 | 0.474 | 0.372 | 0.671 | 0.647 | 0.612 | 0.479 |
|  | sniffles | 0.934 | 0.939 | 0.713 | 0.415 | 0.257 | 0.352 | 0.457 | 0.419 | 0.404 | 0.512 | 0.557 | 0.417 |
| Insertions | dysgu | 0.857 | 0.902 | 0.970 | 0.917 | 0.541 | 0.546 | 0.395 | 0.217 | 0.663 | 0.680 | 0.562 | 0.351 |
|  | nanovar | 0.846 | 0.879 | 0.870 | 0.296 | 0.518 | 0.531 | 0.511 | 0.542 | 0.643 | 0.662 | 0.644 | 0.383 |
|  | svim | 0.826 | 0.863 | 0.964 | 1.000 | 0.555 | 0.518 | 0.350 | 0.079 | 0.664 | 0.647 | 0.514 | 0.147 |
|  | sniffles | 0.872 | 0.905 | 0.962 | 0.750 | 0.434 | 0.475 | 0.301 | 0.142 | 0.579 | 0.623 | 0.459 | 0.239 |

Table S8. 8X PacBio Sequel II reads aligned with ngmlr, All-regions benchmark, split by size.

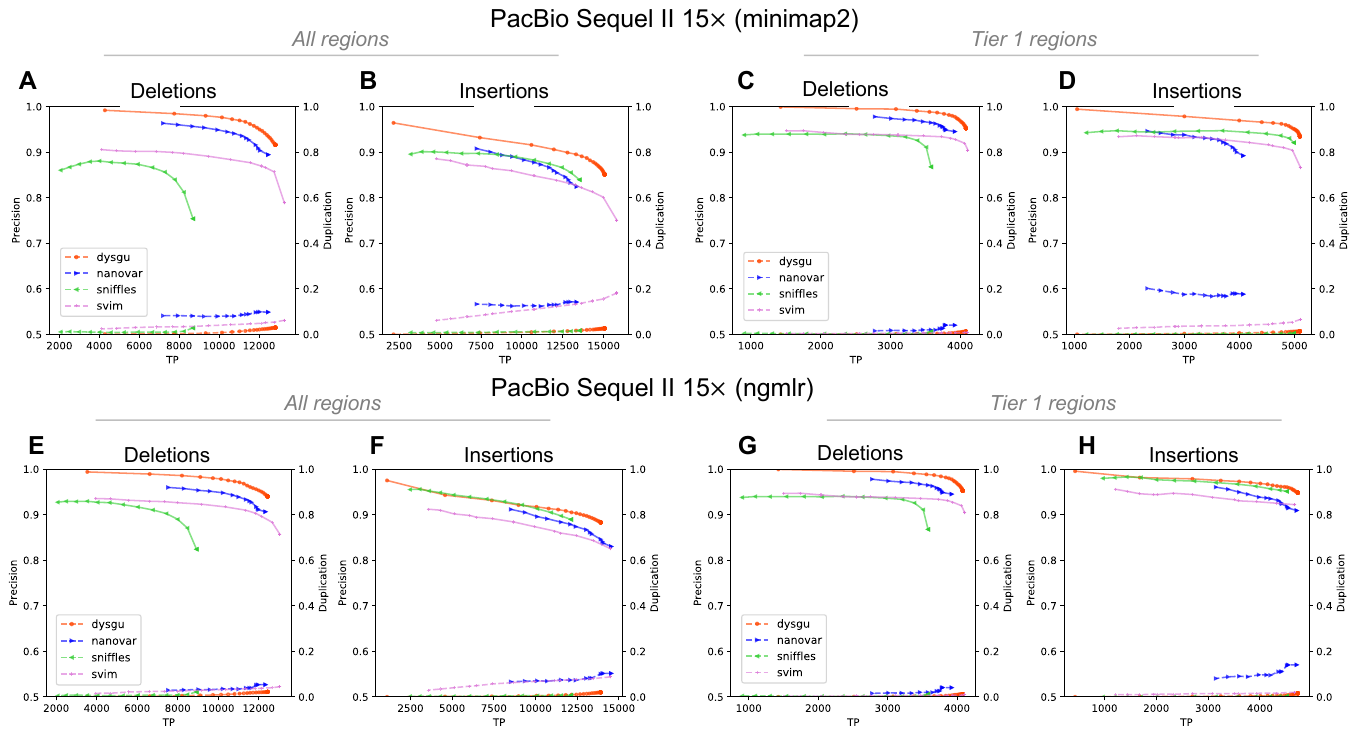

****Figure S3. Performance of dysgu on PacBio reads at 15****$\boldsymbol{\times}$****coverage.****

PacBio Sequel II reads were mapped using minimap2 (A – D) and ngmlr (E – H). Precision-recall curves are shown for deletion and insertion events using the ‘all-regions’ benchmark (A,B,E,F) and Tier 1 high-confidence regions (C,D,G,H).

|  | TP | | FP | | Precision | | Recall | | Duplication | | F1 | |
| --- | --- | --- | --- | --- | --- | --- | --- | --- | --- | --- | --- | --- |
|  | DEL | INS | DEL | INS | DEL | INS | DEL | INS | DEL | INS | DEL | INS |
| dysgu | 4099 | 5172 | 294 | 345 | 0.933 | 0.938 | 0.972 | 0.950 | 0.021 | 0.025 | 0.952 | 0.944 |
| nanovar | 3957 | 4107 | 230 | 344 | 0.945 | 0.923 | 0.938 | 0.755 | 0.129 | 0.202 | 0.942 | 0.830 |
| svim | 4054 | 5143 | 563 | 636 | 0.878 | 0.890 | 0.961 | 0.945 | 0.023 | 0.088 | 0.918 | 0.917 |
| sniffles | 3544 | 5043 | 717 | 363 | 0.832 | 0.933 | 0.840 | 0.927 | 0.029 | 0.008 | 0.836 | 0.930 |

Table S9. 15 X PacBio Sequel II reads aligned using minimap2, Tier1 benchmark.

|  | TP | | FP | | Precision | | Recall | | Duplication | | F1 | |
| --- | --- | --- | --- | --- | --- | --- | --- | --- | --- | --- | --- | --- |
|  | DEL | INS | DEL | INS | DEL | INS | DEL | INS | DEL | INS | DEL | INS |
| dysgu | 4126 | 4899 | 207 | 256 | 0.952 | 0.950 | 0.978 | 0.900 | 0.019 | 0.031 | 0.965 | 0.925 |
| nanovar | 3944 | 4749 | 187 | 385 | 0.955 | 0.925 | 0.935 | 0.873 | 0.045 | 0.150 | 0.945 | 0.898 |
| svim | 4131 | 4700 | 390 | 349 | 0.914 | 0.931 | 0.980 | 0.864 | 0.015 | 0.024 | 0.946 | 0.896 |
| sniffles | 3638 | 4541 | 465 | 206 | 0.887 | 0.957 | 0.863 | 0.834 | 0.020 | 0.004 | 0.875 | 0.891 |

Table S10. 15X PacBio Sequel II reads aligned using ngmlr, Tier1 benchmark.

|  | TP | | FP | | Precision | | Recall | | Duplication | | F1 | |
| --- | --- | --- | --- | --- | --- | --- | --- | --- | --- | --- | --- | --- |
|  | DEL | INS | DEL | INS | DEL | INS | DEL | INS | DEL | INS | DEL | INS |
| dysgu | 12814 | 15156 | 938 | 2284 | 0.932 | 0.869 | 0.343 | 0.415 | 0.034 | 0.042 | 0.501 | 0.561 |
| nanovar | 12602 | 13482 | 1189 | 2320 | 0.914 | 0.853 | 0.337 | 0.369 | 0.108 | 0.169 | 0.492 | 0.515 |
| svim | 13430 | 15932 | 3120 | 4415 | 0.812 | 0.783 | 0.359 | 0.436 | 0.082 | 0.225 | 0.498 | 0.560 |
| sniffles | 9014 | 13728 | 2305 | 2266 | 0.796 | 0.858 | 0.241 | 0.375 | 0.051 | 0.025 | 0.370 | 0.522 |

Table S11. 15X PacBio Sequel II reads aligned using minimap2, All-regions benchmark.

|  | TP | | FP | | Precision | | Recall | | Duplication | | F1 | |
| --- | --- | --- | --- | --- | --- | --- | --- | --- | --- | --- | --- | --- |
|  | DEL | INS | DEL | INS | DEL | INS | DEL | INS | DEL | INS | DEL | INS |
| dysgu | 12873 | 14510 | 859 | 1869 | 0.937 | 0.886 | 0.344 | 0.397 | 0.034 | 0.038 | 0.504 | 0.548 |
| nanovar | 12474 | 14692 | 1044 | 2627 | 0.923 | 0.848 | 0.334 | 0.402 | 0.060 | 0.116 | 0.490 | 0.545 |
| svim | 13203 | 14618 | 1849 | 2687 | 0.877 | 0.845 | 0.353 | 0.400 | 0.058 | 0.103 | 0.504 | 0.543 |
| sniffles | 9184 | 12245 | 1531 | 1338 | 0.857 | 0.902 | 0.246 | 0.335 | 0.034 | 0.012 | 0.382 | 0.488 |

Table S12. 15X PacBio Sequel II reads aligned using ngmlr, All-regions benchmark.

|  |  | Precision | | | | Recall | | | | F1 | | | |
| --- | --- | --- | --- | --- | --- | --- | --- | --- | --- | --- | --- | --- | --- |
|  |  | [30, 50) | [50, 500) | [500, 5000) | ≥5000 | [30, 50) | [50, 500) | [500, 5000) | ≥5000 | [30, 50) | [50, 500) | [500, 5000) | ≥5000 |
| Deletions | dysgu | 0.925 | 0.929 | 0.948 | 0.942 | 0.583 | 0.530 | 0.439 | 0.299 | 0.715 | 0.675 | 0.600 | 0.453 |
|  | nanovar | 0.932 | 0.915 | 0.860 | 0.601 | 0.537 | 0.515 | 0.497 | 0.376 | 0.681 | 0.659 | 0.630 | 0.463 |
|  | svim | 0.834 | 0.778 | 0.830 | 0.919 | 0.604 | 0.553 | 0.483 | 0.301 | 0.701 | 0.647 | 0.611 | 0.453 |
|  | sniffles | 0.903 | 0.847 | 0.611 | 0.272 | 0.310 | 0.407 | 0.478 | 0.389 | 0.461 | 0.550 | 0.536 | 0.320 |
| Insertions | dysgu | 0.819 | 0.859 | 0.958 | 0.921 | 0.605 | 0.619 | 0.532 | 0.229 | 0.696 | 0.719 | 0.684 | 0.367 |
|  | nanovar | 0.835 | 0.848 | 0.851 | 0.287 | 0.534 | 0.512 | 0.554 | 0.206 | 0.651 | 0.639 | 0.671 | 0.240 |
|  | svim | 0.751 | 0.745 | 0.897 | 0.970 | 0.631 | 0.648 | 0.580 | 0.257 | 0.686 | 0.693 | 0.704 | 0.406 |
|  | sniffles | 0.839 | 0.843 | 0.881 | 0.831 | 0.504 | 0.588 | 0.535 | 0.253 | 0.630 | 0.693 | 0.666 | 0.388 |

Table S13. 15X PacBio Sequel II reads aligned with minimap2, All-regions benchmark, split by size.

|  |  | Precision | | | | Recall | | | | F1 | | | |
| --- | --- | --- | --- | --- | --- | --- | --- | --- | --- | --- | --- | --- | --- |
|  |  | [30, 50) | [50, 500) | [500, 5000) | ≥5000 | [30, 50) | [50, 500) | [500, 5000) | ≥5000 | [30, 50) | [50, 500) | [500, 5000) | ≥5000 |
| Deletions | dysgu | 0.945 | 0.964 | 0.961 | 0.837 | 0.544 | 0.506 | 0.456 | 0.262 | 0.690 | 0.664 | 0.619 | 0.399 |
|  | nanovar | 0.934 | 0.933 | 0.894 | 0.516 | 0.520 | 0.504 | 0.497 | 0.389 | 0.668 | 0.654 | 0.639 | 0.443 |
|  | svim | 0.884 | 0.888 | 0.798 | 0.585 | 0.569 | 0.534 | 0.525 | 0.403 | 0.692 | 0.667 | 0.633 | 0.477 |
|  | sniffles | 0.933 | 0.932 | 0.666 | 0.319 | 0.307 | 0.410 | 0.497 | 0.454 | 0.462 | 0.570 | 0.569 | 0.375 |
| Insertions | dysgu | 0.844 | 0.896 | 0.979 | 0.917 | 0.575 | 0.573 | 0.397 | 0.087 | 0.684 | 0.699 | 0.564 | 0.159 |
|  | nanovar | 0.837 | 0.861 | 0.868 | 0.242 | 0.556 | 0.574 | 0.565 | 0.549 | 0.668 | 0.689 | 0.684 | 0.336 |
|  | svim | 0.792 | 0.830 | 0.946 | 1.000 | 0.600 | 0.589 | 0.444 | 0.126 | 0.683 | 0.689 | 0.605 | 0.225 |
|  | sniffles | 0.858 | 0.902 | 0.963 | 0.750 | 0.482 | 0.520 | 0.366 | 0.285 | 0.617 | 0.660 | 0.530 | 0.413 |

Table S14. 15X PacBio Sequel II reads aligned with ngmlr, All-regions benchmark, split by size.

### ONT reads

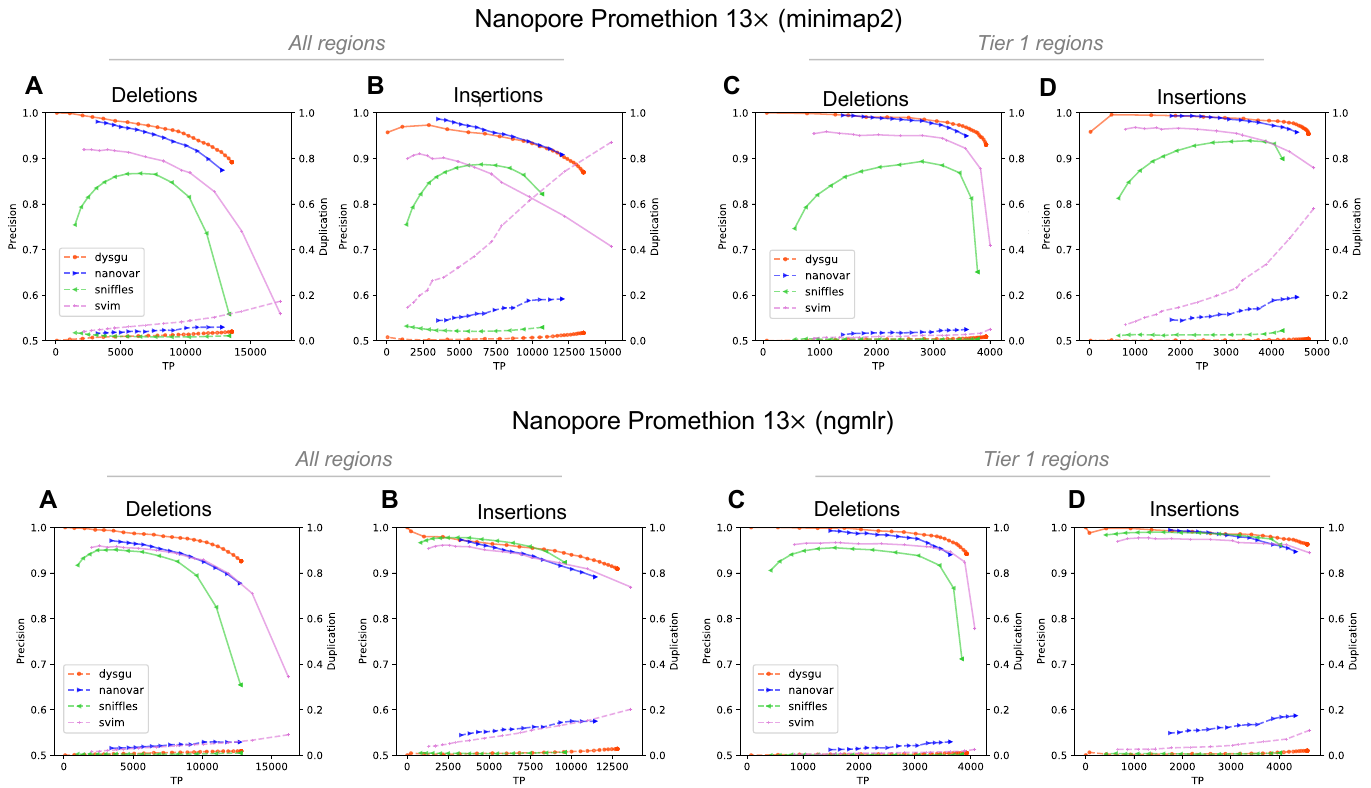

****Figure S4. Performance of dysgu using ONT reads.****

ONT reads were mapped using minimap2 (A – D) and ngmlr (E – H). Precision-recall curves are shown for deletion and insertion events using the ‘all-regions’ benchmark (A,B,E,F) and Tier 1 high-confidence regions (C,D,G,H).

|  | TP | | FP | | Precision | | Recall | | Duplication | | F1 | |
| --- | --- | --- | --- | --- | --- | --- | --- | --- | --- | --- | --- | --- |
|  | DEL | INS | DEL | INS | DEL | INS | DEL | INS | DEL | INS | DEL | INS |
| Dysgu | 3931 | 4811 | 295 | 233 | 0.930 | 0.954 | 0.932 | 0.884 | 0.017 | 0.008 | 0.931 | 0.918 |
| nanovar | 3587 | 4556 | 191 | 202 | 0.949 | 0.958 | 0.851 | 0.837 | 0.049 | 0.191 | 0.897 | 0.893 |
| Svim | 4002 | 4929 | 1638 | 678 | 0.710 | 0.879 | 0.949 | 0.906 | 0.050 | 0.580 | 0.812 | 0.892 |
| sniffles | 3782 | 4238 | 2030 | 475 | 0.651 | 0.899 | 0.897 | 0.779 | 0.007 | 0.044 | 0.754 | 0.835 |

Table S15. 13X ONT Promethion reads aligned using minimap2, Tier1 benchmark.

|  | TP | | FP | | Precision | | Recall | | Duplication | | F1 | |
| --- | --- | --- | --- | --- | --- | --- | --- | --- | --- | --- | --- | --- |
|  | DEL | INS | DEL | INS | DEL | INS | DEL | INS | DEL | INS | DEL | INS |
| dysgu | 3925 | 4579 | 243 | 177 | 0.942 | 0.963 | 0.931 | 0.841 | 0.010 | 0.020 | 0.936 | 0.898 |
| nanovar | 3634 | 4337 | 231 | 244 | 0.940 | 0.947 | 0.862 | 0.797 | 0.059 | 0.173 | 0.899 | 0.865 |
| svim | 4072 | 4628 | 1164 | 274 | 0.778 | 0.944 | 0.966 | 0.850 | 0.026 | 0.110 | 0.862 | 0.895 |
| sniffles | 3841 | 4016 | 1559 | 160 | 0.711 | 0.962 | 0.911 | 0.738 | 0.004 | 0.010 | 0.799 | 0.835 |

Table S16. 13X ONT Promethion reads aligned using ngmlr, Tier1 benchmark.

|  | TP | | FP | | Precision | | Recall | | Duplication | | F1 | |
| --- | --- | --- | --- | --- | --- | --- | --- | --- | --- | --- | --- | --- |
|  | DEL | INS | DEL | INS | DEL | INS | DEL | INS | DEL | INS | DEL | INS |
| dysgu | 13568 | 13506 | 1645 | 2028 | 0.892 | 0.869 | 0.363 | 0.369 | 0.039 | 0.034 | 0.516 | 0.518 |
| nanovar | 12835 | 12051 | 1847 | 1218 | 0.874 | 0.908 | 0.343 | 0.33 | 0.059 | 0.183 | 0.493 | 0.484 |
| svim | 17297 | 15418 | 13642 | 6395 | 0.559 | 0.707 | 0.463 | 0.422 | 0.173 | 0.872 | 0.506 | 0.528 |
| sniffles | 13365 | 10672 | 10571 | 2315 | 0.558 | 0.822 | 0.357 | 0.292 | 0.021 | 0.058 | 0.436 | 0.431 |

Table S17. 13X ONT Promethion reads aligned using minimap2, All-regions benchmark.

|  | TP | | FP | | Precision | | Recall | | Duplication | | F1 | |
| --- | --- | --- | --- | --- | --- | --- | --- | --- | --- | --- | --- | --- |
|  | DEL | INS | DEL | INS | DEL | INS | DEL | INS | DEL | INS | DEL | INS |
| dysgu | 12809 | 12809 | 1023 | 1278 | 0.926 | 0.909 | 0.343 | 0.350 | 0.018 | 0.028 | 0.500 | 0.506 |
| nanovar | 12706 | 11464 | 1774 | 1397 | 0.878 | 0.891 | 0.340 | 0.314 | 0.058 | 0.149 | 0.490 | 0.464 |
| svim | 16207 | 13636 | 7908 | 2055 | 0.672 | 0.869 | 0.434 | 0.373 | 0.090 | 0.202 | 0.527 | 0.522 |
| sniffles | 12759 | 9604 | 6740 | 799 | 0.654 | 0.923 | 0.341 | 0.263 | 0.010 | 0.013 | 0.449 | 0.409 |

Table S18. 13X ONT Promethion reads aligned using ngmlr, All-regions benchmark.

|  |  | Precision | | | | Recall | | | | F1 | | | |
| --- | --- | --- | --- | --- | --- | --- | --- | --- | --- | --- | --- | --- | --- |
|  |  | [30, 50) | [50, 500) | [500, 5000) | ≥5000 | [30, 50) | [50, 500) | [500, 5000) | ≥5000 | [30, 50) | [50, 500) | [500, 5000) | ≥5000 |
| Deletions | dysgu | 0.837 | 0.905 | 0.907 | 0.896 | 0.535 | 0.515 | 0.484 | 0.335 | 0.653 | 0.657 | 0.631 | 0.488 |
|  | nanovar | 0.744 | 0.940 | 0.921 | 0.630 | 0.460 | 0.454 | 0.429 | 0.337 | 0.569 | 0.612 | 0.586 | 0.439 |
|  | svim | 0.342 | 0.670 | 0.823 | 0.913 | 0.622 | 0.569 | 0.499 | 0.323 | 0.442 | 0.616 | 0.621 | 0.477 |
|  | sniffles | 0.354 | 0.662 | 0.708 | 0.425 | 0.461 | 0.463 | 0.487 | 0.397 | 0.400 | 0.545 | 0.577 | 0.410 |
| Insertions | dysgu | 0.841 | 0.849 | 0.942 | 0.936 | 0.464 | 0.595 | 0.538 | 0.522 | 0.598 | 0.700 | 0.685 | 0.670 |
|  | nanovar | 0.932 | 0.901 | 0.887 | 0.602 | 0.358 | 0.544 | 0.562 | 0.455 | 0.517 | 0.678 | 0.688 | 0.518 |
|  | svim | 0.652 | 0.657 | 0.915 | 0.963 | 0.524 | 0.662 | 0.645 | 0.510 | 0.581 | 0.659 | 0.757 | 0.667 |
|  | sniffles | 0.815 | 0.797 | 0.861 | 0.711 | 0.329 | 0.477 | 0.508 | 0.466 | 0.468 | 0.597 | 0.639 | 0.563 |

Table S19. 13X ONT Promethion reads aligned with minimap2, All-regions benchmark, split by size.

|  |  | Precision | | | | Recall | | | | F1 | | | |
| --- | --- | --- | --- | --- | --- | --- | --- | --- | --- | --- | --- | --- | --- |
|  |  | [30, 50) | [50, 500) | [500, 5000) | ≥5000 | [30, 50) | [50, 500) | [500, 5000) | ≥5000 | [30, 50) | [50, 500) | [500, 5000) | ≥5000 |
| Deletions | dysgu | 0.879 | 0.941 | 0.942 | 0.929 | 0.500 | 0.494 | 0.476 | 0.292 | 0.637 | 0.648 | 0.632 | 0.445 |
|  | nanovar | 0.778 | 0.930 | 0.907 | 0.490 | 0.453 | 0.459 | 0.436 | 0.362 | 0.573 | 0.615 | 0.589 | 0.416 |
|  | svim | 0.447 | 0.805 | 0.883 | 0.730 | 0.589 | 0.547 | 0.509 | 0.370 | 0.509 | 0.651 | 0.646 | 0.491 |
|  | sniffles | 0.465 | 0.774 | 0.745 | 0.287 | 0.437 | 0.453 | 0.509 | 0.472 | 0.450 | 0.571 | 0.605 | 0.357 |
| Insertions | dysgu | 0.888 | 0.895 | 0.961 | 1.000 | 0.439 | 0.575 | 0.501 | 0.170 | 0.588 | 0.700 | 0.658 | 0.291 |
|  | nanovar | 0.931 | 0.895 | 0.881 | 0.358 | 0.337 | 0.515 | 0.531 | 0.565 | 0.495 | 0.654 | 0.662 | 0.438 |
|  | svim | 0.818 | 0.858 | 0.958 | 1.000 | 0.475 | 0.609 | 0.493 | 0.174 | 0.601 | 0.712 | 0.651 | 0.296 |
|  | sniffles | 0.903 | 0.920 | 0.945 | 0.810 | 0.300 | 0.441 | 0.412 | 0.372 | 0.451 | 0.596 | 0.573 | 0.509 |

Table S20. 13X ONT Promethion reads aligned with ngmlr, All-regions benchmark, split by size.

### Combinations of sequencing platforms

|  | TP | | Precision | | Recall | | Duplication | | F1 | |
| --- | --- | --- | --- | --- | --- | --- | --- | --- | --- | --- |
|  | DEL | INS | DEL | INS | DEL | INS | DEL | INS | DEL | INS |
| pb 8x | 3869 | 4877 | 0.956 | 0.949 | 0.918 | 0.896 | 0.015 | 0.019 | 0.936 | 0.922 |
| pb 8x + ill 20x | 4004 | 4950 | 0.933 | 0.932 | 0.950 | 0.909 | 0.038 | 0.030 | 0.941 | 0.920 |
| pb 8x + ill 40x | 4018 | 4963 | 0.905 | 0.913 | 0.953 | 0.912 | 0.051 | 0.035 | 0.929 | 0.912 |
| pb 15x | 4056 | 5142 | 0.952 | 0.947 | 0.962 | 0.945 | 0.017 | 0.021 | 0.957 | 0.946 |
| pb 15x + ill 20x | 4103 | 5175 | 0.928 | 0.931 | 0.973 | 0.951 | 0.041 | 0.033 | 0.950 | 0.941 |
| pb 15x + ill 40x | 4103 | 5171 | 0.901 | 0.913 | 0.973 | 0.950 | 0.055 | 0.039 | 0.936 | 0.931 |
| ont 13x | 3931 | 4811 | 0.930 | 0.954 | 0.932 | 0.884 | 0.017 | 0.008 | 0.931 | 0.918 |
| ont 13x + ill 20x | 4016 | 4901 | 0.894 | 0.912 | 0.952 | 0.900 | 0.033 | 0.102 | 0.922 | 0.906 |
| ont 13x + ill 40x | 4033 | 4974 | 0.854 | 0.851 | 0.956 | 0.914 | 0.040 | 0.147 | 0.902 | 0.881 |
| ont 13x + pb 8x | 4105 | 5066 | 0.904 | 0.926 | 0.973 | 0.931 | 0.054 | 0.187 | 0.937 | 0.928 |

Table S21. Combinations of sequencing platforms, Tier1 benchmark.

Calls were made using dysgu on each dataset before merging of outputs. Calls were tested using Tier1 regions on the HG002 benchmark. The coverage value of each sequencing dataset is denoted using ‘x’. pb – PacBio Sequel II, ill – Illumina 150bp PE, ont – Oxford Nanopore Technologies Promethion.

|  | TP | | Precision | | Recall | | Duplication | | F1 | |
| --- | --- | --- | --- | --- | --- | --- | --- | --- | --- | --- |
|  | DEL | INS | DEL | INS | DEL | INS | DEL | INS | DEL | INS |
| pb 8x | 8587 | 10671 | 0.952 | 0.921 | 0.442 | 0.501 | 0.045 | 0.047 | 0.604 | 0.649 |
| pb 8x + ill 20x | 9076 | 10988 | 0.948 | 0.920 | 0.467 | 0.516 | 0.063 | 0.060 | 0.626 | 0.661 |
| pb 8x + ill 40x | 9261 | 11173 | 0.944 | 0.918 | 0.477 | 0.525 | 0.074 | 0.066 | 0.634 | 0.668 |
| pb 15x | 9002 | 11255 | 0.952 | 0.920 | 0.464 | 0.528 | 0.047 | 0.049 | 0.624 | 0.671 |
| pb 15x + ill 20x | 9343 | 11483 | 0.948 | 0.919 | 0.481 | 0.539 | 0.065 | 0.062 | 0.638 | 0.679 |
| pb 15x + ill 40x | 9482 | 11598 | 0.945 | 0.917 | 0.488 | 0.545 | 0.078 | 0.067 | 0.644 | 0.683 |
| ont 13x | 9139 | 10657 | 0.948 | 0.914 | 0.471 | 0.500 | 0.054 | 0.038 | 0.629 | 0.647 |
| ont 13x + ill 20x | 9667 | 11191 | 0.939 | 0.910 | 0.498 | 0.525 | 0.072 | 0.118 | 0.651 | 0.666 |
| ont 13x + ill 40x | 9905 | 11488 | 0.934 | 0.904 | 0.510 | 0.539 | 0.083 | 0.150 | 0.660 | 0.676 |
| ont 13x + pb 8x | 10030 | 11863 | 0.913 | 0.880 | 0.516 | 0.557 | 0.129 | 0.200 | 0.660 | 0.682 |

Table S22. Combinations of sequencing platforms, Tier1+2 benchmark.

Calls were made using dysgu on each dataset before merging of outputs. Calls were tested using Tier1+2 regions on the HG002 benchmark. The coverage value of each sequencing dataset is denoted using ‘x’. pb – PacBio Sequel II, ill – Illumina 150bp PE, ont – Oxford Nanopore Technologies Promethion.
